# All-optical electrophysiology reveals behavior-dependent dynamics of excitation and inhibition in the hippocampus

**DOI:** 10.1101/2025.03.20.644347

**Authors:** Qixin Yang, Shulamit Baror-Sebban, Rotem Kipper, Michael London, Yoav Adam

## Abstract

Understanding how neuronal integration is modulated by behavior is a fundamental goal in neuroscience. We combined voltage imaging with optogenetic depolarization to reveal how changes in excitatory (E) and inhibitory (I) inputs, modulate the spiking output, subthreshold dynamics, and gain of key genetically defined cell types in the CA1 region of the hippocampus. We imaged pyramidal cells (PCs), vasoactive intestinal peptide (VIP), somatostatin (SST), and parvalbumin (PV) interneurons (INs) and found that locomotion reduced firing in PCs and VIP INs while increasing activity in SST and PV cells. Prolonged optical depolarization experiments and simulations revealed that intracellular theta oscillations are predominantly driven by inhibitory inputs in PCs and VIP cells. Firing rate-laser intensity (F-I) curves revealed distinct gain modulation across cell types, with a divisive gain reduction in PC bursting during locomotion, while simple spikes are unaffected. A two-compartment model suggested that this results from a balanced E/I input increase to somatic and dendritic compartments. These findings reveal how behavioral state-dependent coordination of excitation and inhibition governs hippocampal neuronal dynamics and output-specific gain modulation.

## Introduction

Neuronal integration is a fundamental process by which neurons combine multiple excitatory and inhibitory synaptic inputs to modulate subthreshold dynamics and generate spiking outputs^1–3^. Sensory inputs and behavioral states dynamically influence neuronal integration, leading to gain modulation—a key mechanism that adjusts neuronal responsiveness to changing inputs and ensures efficient information processing^4–7^. Locomotion-dependent changes in gain highlight the context-specific nature of these processes across brain regions. In the primary visual cortex, locomotion enhances gain through increased depolarization and synaptic fluctuations, amplifying responses to visual stimuli and improving signal-to-noise ratios^5,8^. In contrast, in the primary auditory cortex, locomotion reduces gain through mechanisms mediated by local INs^9^, reflecting the region-specific modulation of sensory inputs based on behavioral demands. These examples demonstrate how gain modulation dynamically optimizes sensory processing in response to changing behavioral states.

While gain modulation has been extensively studied in sensory cortices, its exploration in regions involved in higher cognitive functions such as the hippocampus, remains challenging. The hippocampus plays a central role in cognitive processes such as navigation, spatial learning, and memory^10^. The CA1 region, in particular, has a well-defined microcircuitry that parallels other cortical regions^11^. CA1 PCs, the principal excitatory output neurons, receive excitatory inputs both directly via the perforant path from layer III of the entorhinal cortex (EC3) and indirectly via the Schaffer collaterals from CA3 PCs^12^. In addition, PCs are tightly regulated by several types of local inhibitory INs, including VIP, PV, and SST INs (Figure 2A). Together, these excitatory and inhibitory components contribute to the fine-tuning of the E/I balance, allowing the circuit to dynamically adjust its activity in response to changes in behavioral states^13^.

Understanding neuronal integration within the CA1 microcircuit requires access to both subthreshold inputs and spiking outputs. Investigating gain modulation in this region is further complicated by the fact that hippocampal neurons are not tuned to single sensory modalities, unlike neurons in sensory cortices where graded input strength can be manipulated via sensory stimuli^8^. In CA1, it is thus necessary to directly control the input strength, posing a unique technical challenge for dissecting gain modulation mechanisms *in vivo*. Traditional approaches, such as *in vivo* calcium imaging and extracellular electrophysiology, offer important insights but either provide an indirect measurement of spiking activity (calcium Imaging)^14^ or lack the ability to target single cells for manipulation (extracellular electrophysiology). Importantly, neither technique captures the subthreshold membrane potential changes essential for understanding how neurons process synaptic inputs. *In vivo* patch-clamp recordings provide high temporal resolution and access to subthreshold dynamics but are technically challenging to perform, particularly in behaving animals^15–19^.

To address these limitations, we utilized voltage imaging combined with optogenetic depolarization of targeted single cells to study how the CA1 microcircuit integrates E/I inputs during behavior. Voltage imaging provides a direct, high-resolution readout of membrane potential, capturing both spiking and subthreshold activity across multiple genetically defined cell types^20^. Near-infrared GEVIs based on Archaerhdopsin are optically orthogonal to blue-shifted channelrhodopsins and thus allow for precise control and readout of neuronal responses when co-expressed together^21,22^. This combination offers a unique opportunity to evaluate input-output transformations in the neuronal activity of various cell types during behavior^20,23–26^.

In previous work, we developed a set of protocols utilizing an all-optical electrophysiology approach to investigate how CA1 SST INs integrate E/I inputs in awake, behaving mice^23^. Here we expanded these studies to comprehensively examine the activity of key genetically defined CA1 cell types, including PCs which are the main hippocampal output, as well as various local IN types, during two behavioral states - quiet and locomotion. Our analysis focused on changes in firing patterns, subthreshold oscillations, and gain modulation. We revealed how gain modulation acts only on a specific type of PC output pattern, namely spike burst, and using computational modeling we show that increased activity of both soma and dendrite targeting INs during locomotion could provide a mechanism for this phenomenon. Finally, we developed a classifier to distinguish cell types based on intracellular spike waveforms. Overall, our study offers an in-depth view of the intricate interactions among hippocampal neurons that drive state-dependent regulation of neuronal activity, theta oscillations, and E/I balance. Furthermore, it highlights new opportunities opened by voltage-imaging-based all-optical electrophysiology in dissecting neuronal circuit function at unprecedented resolution and detail.

## Results

### Cell-type specific all-optical electrophysiology

We expressed an Optopatch construct^21,23^ composed of a near-infrared (NIR) genetically encoded voltage indicator (GEVI) and the blue-shifted channelrhodopsin somCheRiff in 4 major CA1 cell types: VIP, SST, and PV inhibitory INs, as well as in excitatory PCs (Figures 1A and 1C). We used the GEVI somArchon1^26^ in all cell types, except for fast-spiking PV INs in which we expressed the faster GEVI QuasAr6b^27^. Our approach for high signal-to-noise ratio (SNR) one-photon voltage imaging relies on the sparse expression of the GEVI combined with targeted illumination^23^. INs spatial distribution is naturally sparse, thus using Cre-dependent AAV vectors in combination with standard Cre driver lines (VIP-Cre, SST-Cre, and PV-Cre) was sufficient (Figures 2A-2C). CA1 PCs are densely packed and to control for the sparseness we used an intersectional approach. Specifically, we injected CamKII-Cre mice, which label all PCs, with a Flp-dependent Optopatch vector mixed with a low titer Cre-dependent Flp virus (Figure S2D, see ‘Methods’ section for details).

**Figure 1.**
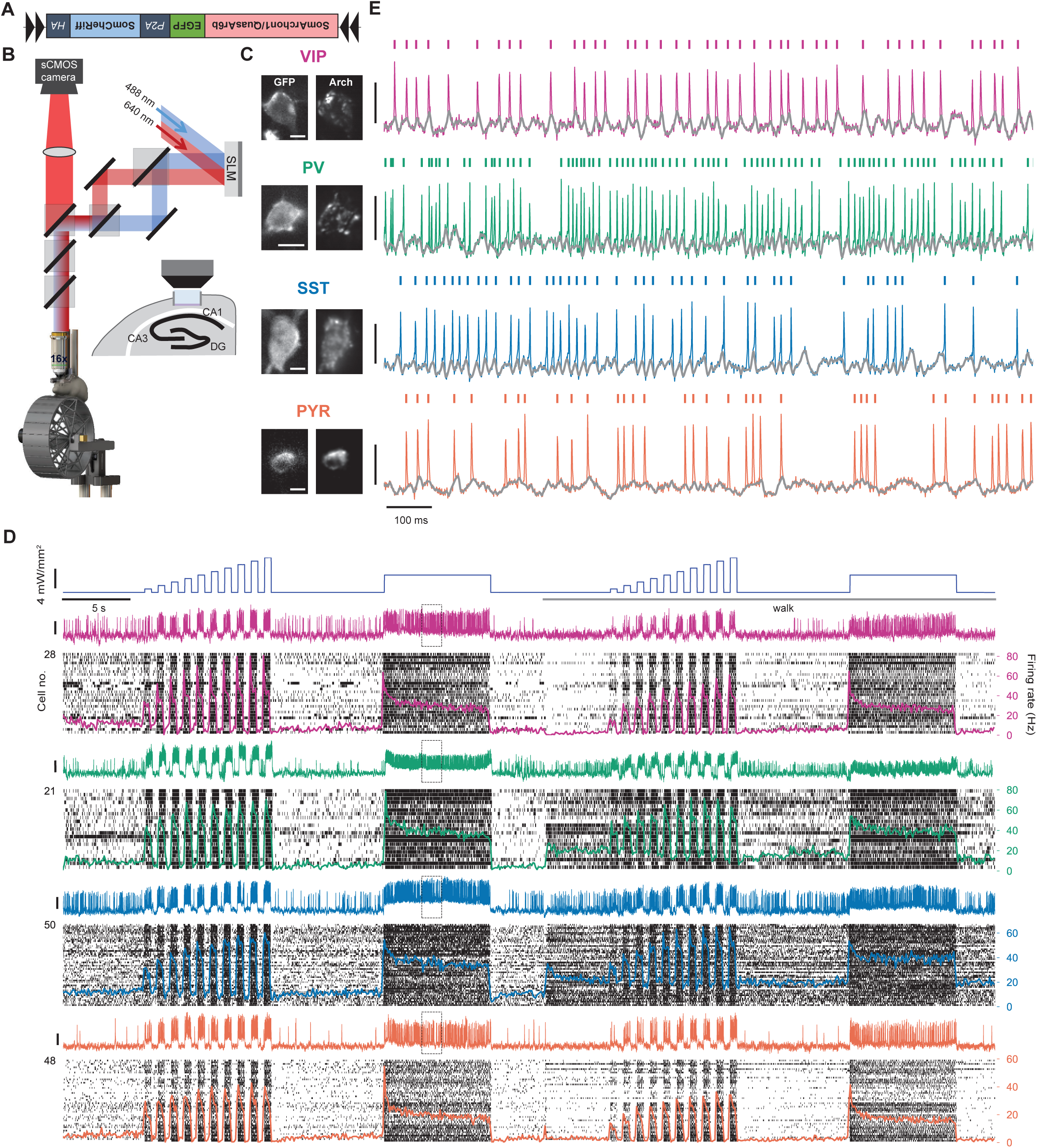
Cell-type specific voltage imaging and optogenetic stimulation in CA1 neurons. (A) Schematic of Cre-dependent viral constructs used for co-expression of the genetically encoded voltage indicator (SomArchon1 for SST, VIP, and CKII-Cre mice, and QuasAr6b for PV-Cre mice) and the optogenetic actuator SomCheriff. (B) Optical setup for simultaneous voltage imaging and optogenetics using red (640 nm) and blue (488 nm) lasers. The system includes dual-channel patterned illumination through a spatial light modulator (SLM). (C) Left, epifluorescence images with wide-field blue illumination of GFP. Right, the same field of view with patterned red illumination of SomArchon1 (QuasAr6b for PV interneurons). Scale bars, 10 µm. (D) Top, optogenetic stimulation protocol during quiet and walking. Bottom, colored traces represent example fluorescence recordings for each cell type. Raster plots display spiking activity from 28 VIP INs, 21 PV INs, 50 SST INs, and 48 PCs, N=3 mice for each cell type with population-averaged firing rate overlaid on top. Vertical scale bars, SNR = 5. (E) Example fluorescence recordings from fields of view shown in (C), enlarged from traces shown in (D). Colored ticks denote identified spikes, and gray lines represent subthreshold activity after spike removal. Vertical scale bar, SNR = 5.

Targeted illumination was achieved using a custom imaging system equipped with a spatial light modulator (SLM) that independently patterned a red 640 nm and a blue 488 nm laser (Figure 1B). Cells were identified using widefield imaging of the GFP tag attached to the GEVI (Figure S2) and a manual region of interest was drawn around selected cells and was converted to phase masks that targeted both the red and blue lasers to illuminate the selected cells. Fluorescence was recorded using continuous 640 nm illumination at the intensity of 15 W/mm^2^ and collected onto a scientific CMOS camera at 1000 frames/sec.

To study brain-state-dependent changes in neuronal activity we recorded from head-fixed mice whose locomotion was controlled with a motorized treadmill. Each cell was recorded during 35 seconds of rest followed by 35 seconds of controlled walking at the speed of 10 cm/s. The optogenetic stimulation protocol included both continuous 8-second illumination at 4 mW/mm^2^ and 10 steps of 500 ms on 500 off at linearly increased intensities from 0.8 to 8 mW/mm^2^. Importantly, the optogenetic stimulation was always targeted to a single cell within an FOV. This protocol was repeated during the resting and walking epochs (Figure 1D, top). We analyzed only cells that showed spiking activity, and all these cells responded to the optogenetic depolarization with increased firing rates (Figure 1D). The imaging SNR allowed high-fidelity detection of action potentials as well as the subthreshold activity (Figure 1E) and thus provided us with a comprehensive characterization of the input-output activity of all cells.

### Behavioral state-dependent firing of CA1 neurons supports the canonical microcircuit organization

The CA1 microcircuitry shows the canonical cortical microcircuit organization, with PCs as the primary excitatory output, intricately regulated by dendritic-targeting SST INs and perisomatic-targeting PV INs^11^. Additionally, VIP-expressing INs serve a disinhibitory role by inhibiting the INs that suppress PCs (Figure 2A).

**Figure 2.**
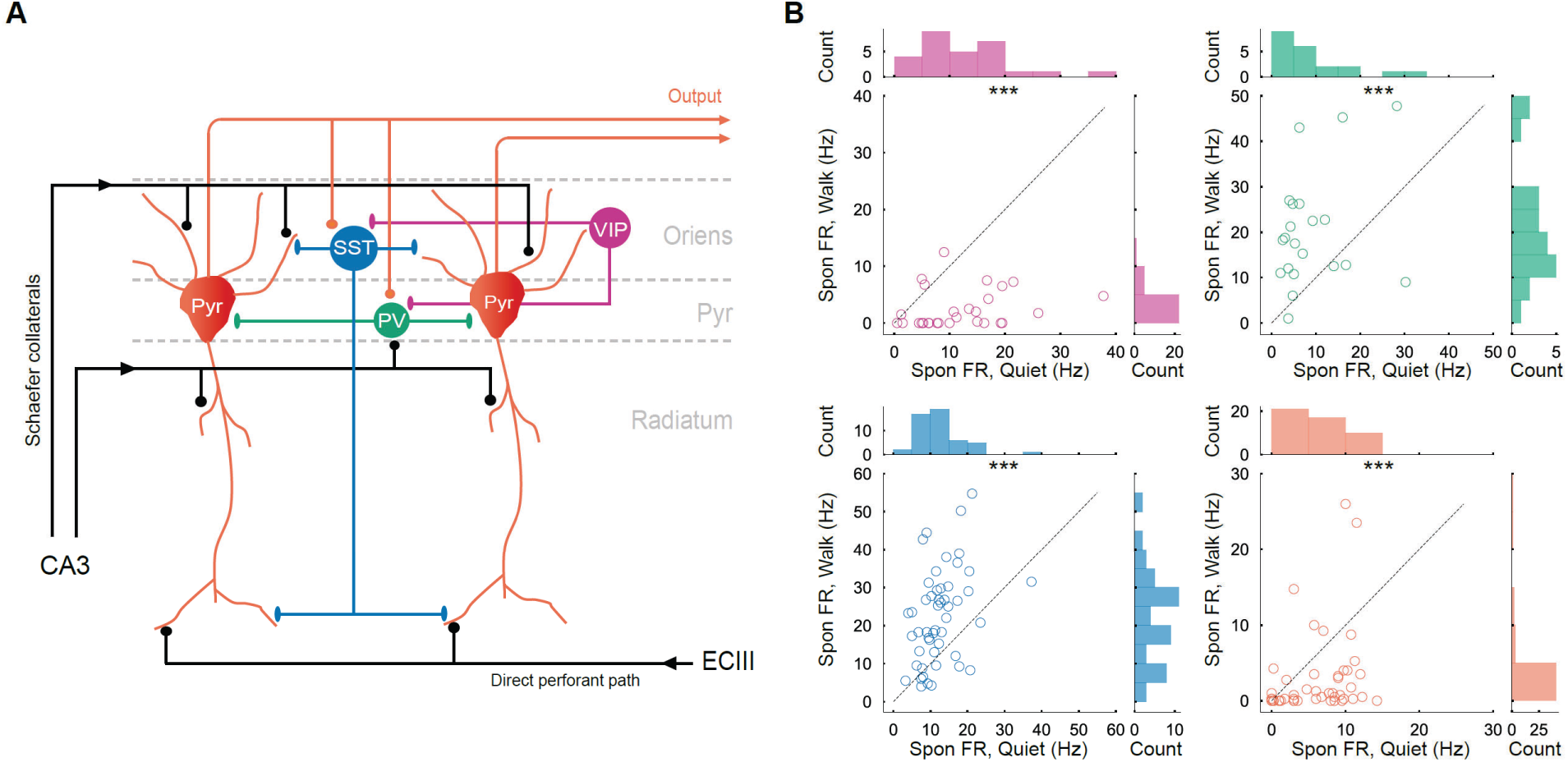
Behavioral state-dependent firing in PCs and identified INs in CA1. (A) Schematic representation of the CA1 microcircuit, illustrating the interactions between PCs and various types of INs. (B) Spontaneous firing rates of VIP, PV, SST INs, and PCs during the 4 seconds preceding the F-I protocol in quiet and walking conditions. Top row, histograms and scatter plots showing the spontaneous firing rates of VIP and PV INs. Bottom row, histograms and scatter plots showing the spontaneous firing rates of SST INs and PCs. Each scatter plot compares spontaneous firing rates between quiet and walking conditions. Statistical significance is denoted by *** (p < 0.001).

We analyzed the firing rates of all cell types and found that VIP INs and PCs exhibited decreased firing rates during locomotion compared to quiet periods (VIP: quiet, 11.9 ± 1.6 Hz (mean ± s.e.m.); walk, 2.4 ± 0.6 Hz; p = 1.8×10^-6^, paired t-test. PC: quiet, 5.8 ± 0.6 Hz; walk, 2.9 ± 0.8 Hz; p = 3.0×10^-4^, Wilcoxon signed-rank test. Figure 2B). In contrast, PV and SST INs showed increased firing rates during walking (PV: quiet, 8.5 ± 1.8 Hz; walk, 19.9 ± 2.5 Hz; p = 1.6×10^-5^, paired t-test. SST: quiet, 12.5 ± 0.8 Hz; walk, 22.5 ± 1.7 Hz; p = 1.1×10^-7^, paired t-test; Figure 2B). This pattern is in agreement with the established CA1 microcircuit model (Figure 2A), wherein reduced VIP IN activity during walking leads to increased activity of SST and PV INs, thereby enhancing their inhibitory influence on PCs and reducing PC activity. This model primarily considers local connectivity and does not account for alterations in long-range inputs to PCs, such as those from CA3 and EC3 regions. However, our results emphasize an enhanced inhibitory control of CA1 output during locomotion, which implies strengthening of feedforward or feedback inhibition within the CA1 microcircuit^28–30^. Such behavioral state-dependent inhibitory control is beneficial as it could amplify the spatial selectivity of PCs and promote sparse coding, enhancing the efficiency of information processing within the hippocampus^30,31^.

### Cell type-specific inputs drive intracellular theta oscillations

A key advantage of voltage imaging is that it provides access not only to the spiking dynamics but also to the underlying subthreshold activities. Theta oscillations, rhythmic fluctuations in neural activity within the 4-12 Hz range, play an important role in spatial navigation, learning, and memory^10^. Traditionally recorded as local field potentials, these oscillations reflect population-level phenomena driven by local and long-range E/I synaptic inputs^10^. While intracellular theta oscillations have been documented in CA1 PCs and putative INs using *in vivo* whole-cell recordings ^15–17,32–34^ and voltage imaging^23,26,35^, how these oscillations are modulated by behavioral states at the single-cell level, as well as the inputs driving these modulations, remains unclear.

To address these questions, we examined membrane potential oscillations at the theta frequency band in all 4 cell types. Our results revealed strong intracellular theta oscillations in VIP, PV, and SST INs during walking (Figure 3A and 3B). As we reported previously^23^, locomotion reduced the low-frequency oscillations in CA1 PCs, resulting in a clear peak at 8 Hz which was much weaker compared with all the IN subtypes (Figure 3B). As expected, intracellular theta oscillations were absent across all cell types during periods of quiet, aligning with the established understanding that theta oscillations predominantly emerge during exploratory behaviors^10^.

**Figure 3.**
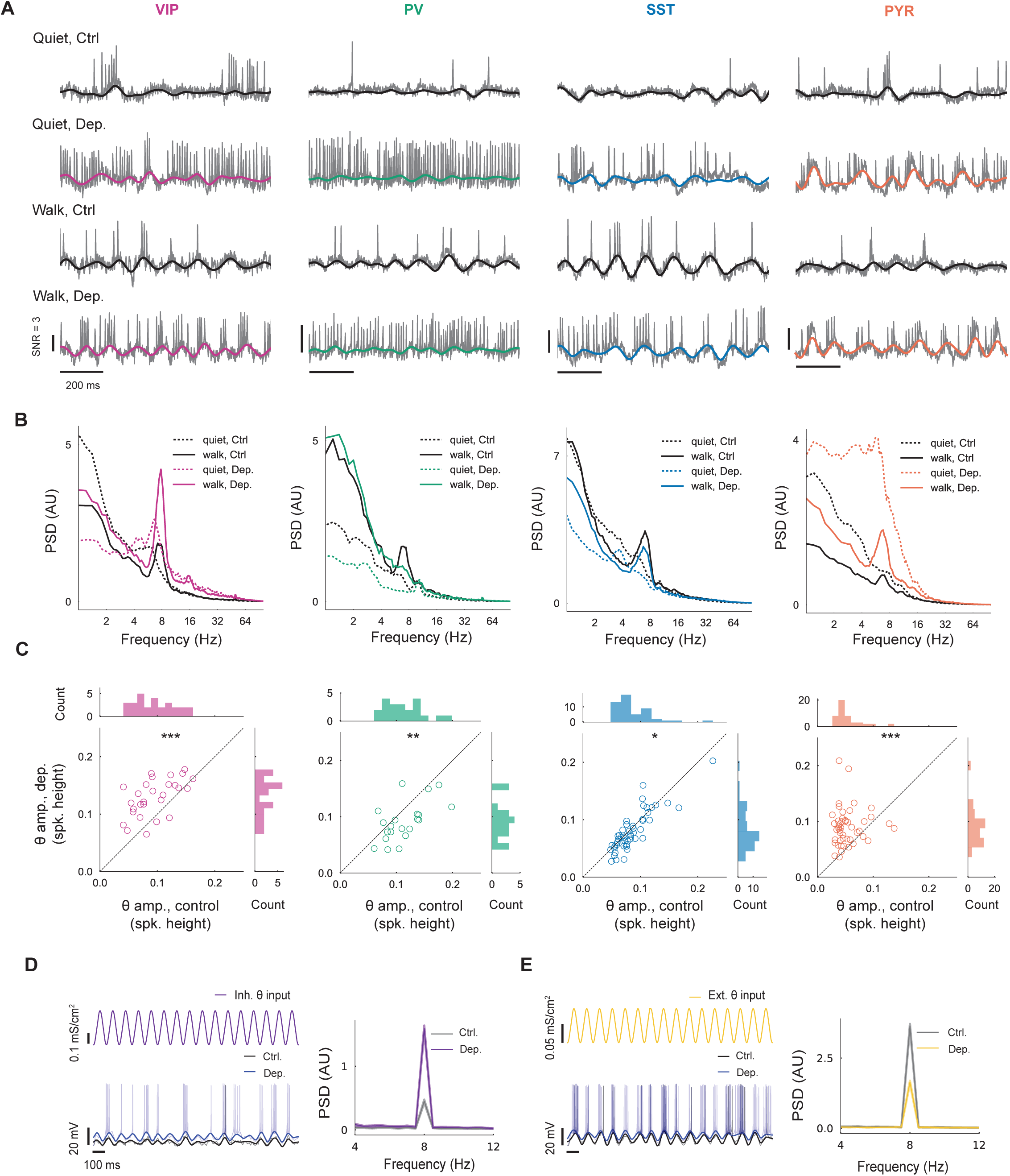
Intracellular theta oscillations in CA1 neurons are modulated by behavioral states and optogenetic stimulation. (A) Representative traces of VIP, PV, SST INs, and PCs with or without optogenetic stimulation, during quiet and walking states. Gray, raw fluorescence. Black and colored, subthreshold 8 Vm (spike removed, bandpass filtered between 4 and 12 Hz). (B) Effects of behavior state and optogenetic stimulation on population averaged PSDs. (C) Theta amplitude of individual cell with or without optogenetic stimulation during walking, calculated as the mean amplitude of Hilbert transformed 8 Vm (6-9 Hz). Significance: * p < 0.05, ** p < 0.01, *** p < 0.001 (paired t-test or Wilcoxon signed-rank test). (D) Left, example membrane voltage traces of a modeled neuron receiving only inhibitory input oscillating at 8 Hz, with or without prolonged depolarization. Right, effect of prolonged depolarization on the mean PSD of the modeled neuron (n=50 simulations). (E) Same as (D), but for a modeled neuron receiving excitatory input oscillating at 8 Hz.

To reveal the contributions of excitatory and inhibitory oscillating inputs to membrane potential theta oscillations during walking, we applied an 8-second constant optogenetic stimulation at 4 mW/mm² during both behavioral states^23^. This depolarization was employed to enhance the inhibitory driving force while reducing the excitatory driving force^23^. During walking, depolarization significantly enhanced intracellular theta oscillations in VIP INs and PCs (VIP, p = 3.2×10^-6^, paired t-test; PC, p = 1.5×10^-8^, Wilcoxon signed-rank test; Figure 3B and 3C). In contrast, PV cells demonstrated a substantial reduction in their mean intracellular theta amplitude, and SST INs exhibited a comparatively modest decrease. (PV, p = 0.0029; SST, p = 0.025, paired t-test; Figure 3B and 3C).

To interpret these observations, we performed simulations using a single-compartment conductance-based model (based on the Hodgkin-Huxley model) to assess neuronal responses to E/I theta-oscillating synaptic inputs, with or without prolonged depolarization. The model predicted that membrane potential theta oscillations amplitude is enhanced during depolarization for a purely inhibitory input, and suppressed when the input is excitatory (Figure 3D and 3E). For the more realistic case of mixed E/I inputs with varying strengths and phase differences, the simulations showed that a sufficiently higher inhibitory theta input compared to excitatory theta input (an I/E ratio above 4) is required to enhance intracellular theta oscillations under constant depolarization, regardless of phase differences (Figure S4E). However, at an I/E ratio above 1 but below 4, theta suppression may occur, particularly when the phase difference is small. Importantly, overwhelming excitatory input (I/E<1) consistently induced strong suppression of intracellular theta oscillations.

Based on these simulations, we infer that PCs and VIP INs predominantly receive inhibitory theta-modulated input during walking, whereas PV and SST INs are likely more influenced by excitatory inputs. However, it remains unclear whether the I/E ratio in these cells is less than 1, a condition where excitation would dominate. Previous *in vitro* whole-cell recordings have demonstrated that CA1 PCs receive inhibitory postsynaptic potentials, whereas fast-spiking INs receive excitatory postsynaptic potentials during spontaneous theta oscillations^36^. These findings align with our observations for PCs and provide further evidence that PV INs predominantly receive excitatory theta-modulated inputs.

Furthermore, we observed that PV and SST INs spike rhythmically at a theta frequency during walking (Figure S3A), indicating that they could serve as potential local sources of theta-oscillating inhibitory input to the somatic and dendritic compartments of PCs, respectively. Combined with previous findings that theta oscillations can be self-generated within CA1 without extrinsic long-range inputs^36^, this suggests that these local INs may play a crucial role in sustaining intrinsic theta rhythms within the CA1 microcircuit. PCs firing seemed weakly theta modulated (Figure S3A), possibly due to low firing rates. However, when additional external input was provided using optogenetic stimulation, theta-modulated firing strengthened in the PCs (Figure S3B), further suggesting that PCs are subjected to strong local inhibitory control during walking.

### Different modes of gain modulation

Neuronal gain is critically modulated by changes in E/I balance^37^, but this property is hard to measure during behavior, particularly in the case of hippocampal circuits which are not driven by scaled sensory inputs. We used an all-optical approach to directly record neuronal gain across different E and I cell types and behavioral states. Specifically, we measured the firing rate response to 10 linearly increasing optogenetic stimulation strengths during both quiet and walking conditions, resulting in firing rate versus laser intensity curves (i.e., optical F-I curves)^23^ (Figures 4A-4C). The optogenetic stimulations were 500 ms in duration, separated by 500 ms intervals, with intensities ranging from 0.8 to 8 mW/mm². Given that many cells exhibited response saturation (Figure 4B) and some PCs demonstrated a high response threshold below which no firing occurs (Figure S5), we used a sigmoidal function to fit the F-I curves (27 out of 29 VIP INs, 20 out of 22 PV INs, 46 out of 50 SST INs, 45 out of 48 PCs showed goodness of fit greater than 0.1 in both quiet and walking conditions and were included in the subsequent analysis). The F-I slope was then calculated from the most linear segment of the fitted F-I curve (see “Methods”).

**Figure 4.**
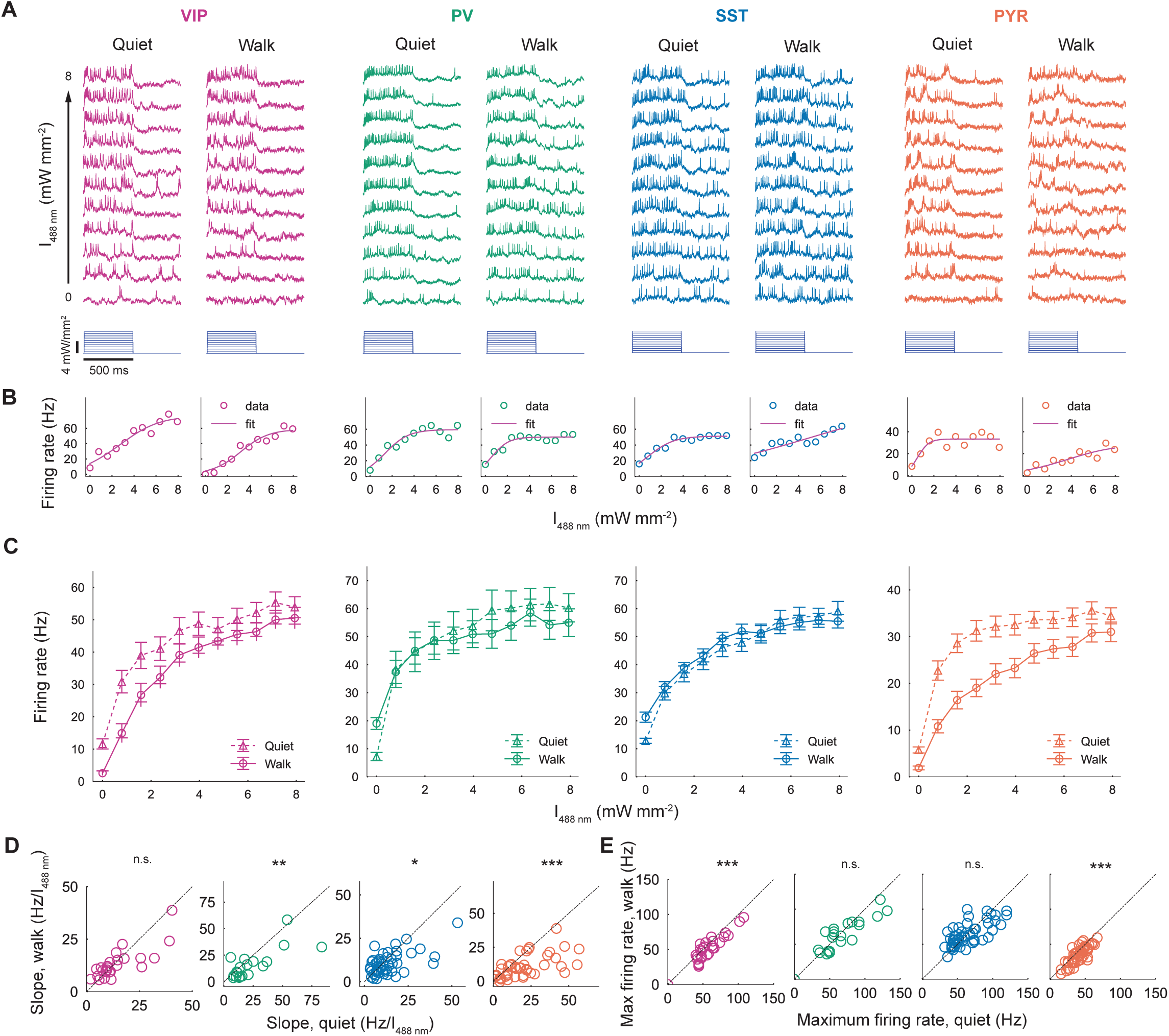
Walking evokes differential gain modulation in CA1 neurons. (A) Fluorescence response of single neurons from the indicated populations to 500-ms optogenetic stimulation steps ranging from 0 to 5 mW/mm² during quiet and walking states. (B) Firing rate plotted against the intensity of optogenetic stimulation (F-I curve) during quiet (left) and walking (right). Magenta lines indicate the fitted sigmoidal functions for the F-I curves. (C) Comparison of population-averaged F-I curves during quiet and walking states, illustrating gain modulation for all cell types except VIP INs (data are mean ± s.e.m). (D) Comparison of slopes of the F-I curves between quiet and walking conditions. (E) Comparison of maximum firing rates between quiet and walking conditions.

Our results revealed three distinct modes of behavior-dependent F-I modulation across the examined cell types (Figure 4C). For VIP INs, the average F-I curve exhibited a subtractive shift during walking, with no significant slope change (quiet, 19.0 ± 3.6 Hz/I_488 nm_; walk, 16.6 ± 3.0 Hz/I_488 nm_; p = 0.86, Wilcoxon signed-rank test; Figure 4D), and reduction of the maximum firing rate during walking (quiet, 58.25 ± 3.9 Hz; walk, 51.73 ± 3.6 Hz; p = 7.3×10^-4^, Wilcoxon signed-rank test; Figure 4E). In contrast, SST and PV INs displayed enhanced firing rates only at lower levels of optogenetic stimulation during walking, with a decrease in the F-I slope (PV, quiet, 29.9 ± 7.4 Hz/I_488 nm_; walk, 18.4 ± 3.9 Hz/I488 nm; p = 2.7×10^-3^, Wilcoxon signed-rank test. SST, quiet, 13.7 ± 1.55 Hz/I_488 nm_; walk, 10.5 ± 1.0 Hz/I_488 nm_; p = 0.025, Wilcoxon signed-rank test.) while their maximum firing rate remained unchanged (PV, quiet, 66.1 ± 6.7 Hz; walk, 61.8 ± 5.4 Hz; p=0.17, paired t-test. SST, quiet, 65.1± 3.3 Hz; walk: 64.4 ± 2.4 Hz; p = 0.78, paired t-test). PCs showed pronounced gain reduction (divisive scaling of the F-I curve marked by a change in the slope) during walking compared to quiet, with the F-I slope significantly decreasing during walking (quiet, 19.9 ± 2.3 Hz/I_488 nm_; walk, 10.4 ± 1.3 Hz/I_488 nm_; p = 4.9×10^-7^, Wilcoxon signed-rank test), as well as the maximum firing rate (quiet, 40.6 ± 1.8 Hz; walk: 34.5 ± 1.8 Hz; p = 2.3×10^-6^, paired t-test).

These *in vivo* findings align with mechanisms previously described in brain slice dynamic patch-clamp studies that simulated *in vivo*-like synaptic inputs^37^. Based on these findings, the three observed patterns of F-I curve modulation can be attributed to distinct changes in E/I synaptic input configurations. A subtractive shift in the F-I curve, as seen in VIP INs, is consistent with increased shunting (i.e., changes in conductance without affecting input variance). The enhancement in neuronal response at low firing rates, observed in SST and PV INs, aligns with an increase in input variance while maintaining constant mean levels of excitatory and inhibitory conductance. Finally, the divisive gain reduction observed in PCs potentially corresponds to a balanced increase in both excitatory and inhibitory background firing rates, reflecting a combined effect of changes in conductance and variance in background inputs^37^.

### Gain modulation in Pyramidal cells is specific for bursts

CA1 PCs are prone to bursts, which are groups of closely timed spikes. These bursts play a key role in information transmission and synaptic plasticity^15,38,39^. Previous intracellular studies have shown that these bursts are typically accompanied by complex subthreshold features, including a large subthreshold depolarization envelope (DE) and after-depolarization potentials (ADPs), also known as plateau potentials ^15,16,34,40,41^. We utilized both an inter-spike interval (ISI) threshold and a depolarization threshold to distinguish bursts from simple spikes^34^ (see “Methods”; Figure 5A, Figure S6A). We found that the ISI between the first and second spike remained consistent across states (quiet: 7.7 ± 0.18 ms; walk: 8.1 ± 0.37 ms; p = 0.79, Mann-Whitney U test), aligning with previous intracellular studies^16,34^. However, we observed clear distinction in burst waveforms across the two behavioral states (Figures 5B and S6B). Interestingly, re-analysis of F-I curves after event-type classification revealed minimal gain changes for simple spikes (Figure 5B, top), whereas burst event rates exhibited significant divisive gain reduction during walking (Figure 5B, bottom) as the slopes of F-I curves for bursts significantly decreased (quiet, 2.8 ± 0.3 Hz/I_488 nm_ ; walk, 1.6± 0.3 Hz/I_488 nm_; p=3.1×10^-6^, paired t-test), while those for simple spikes remained unchanged (quiet, 5.1 ± 0.7 Hz/I_488 nm_ ; walk, 4.0 ± 0.6 Hz/I_488 nm_; p=0.15, paired t-test; Figure 5D). These results suggest that strong gain modulation in PCs (Figures 4C-4E) is exclusively associated with bursting output.

**Figure 5.**
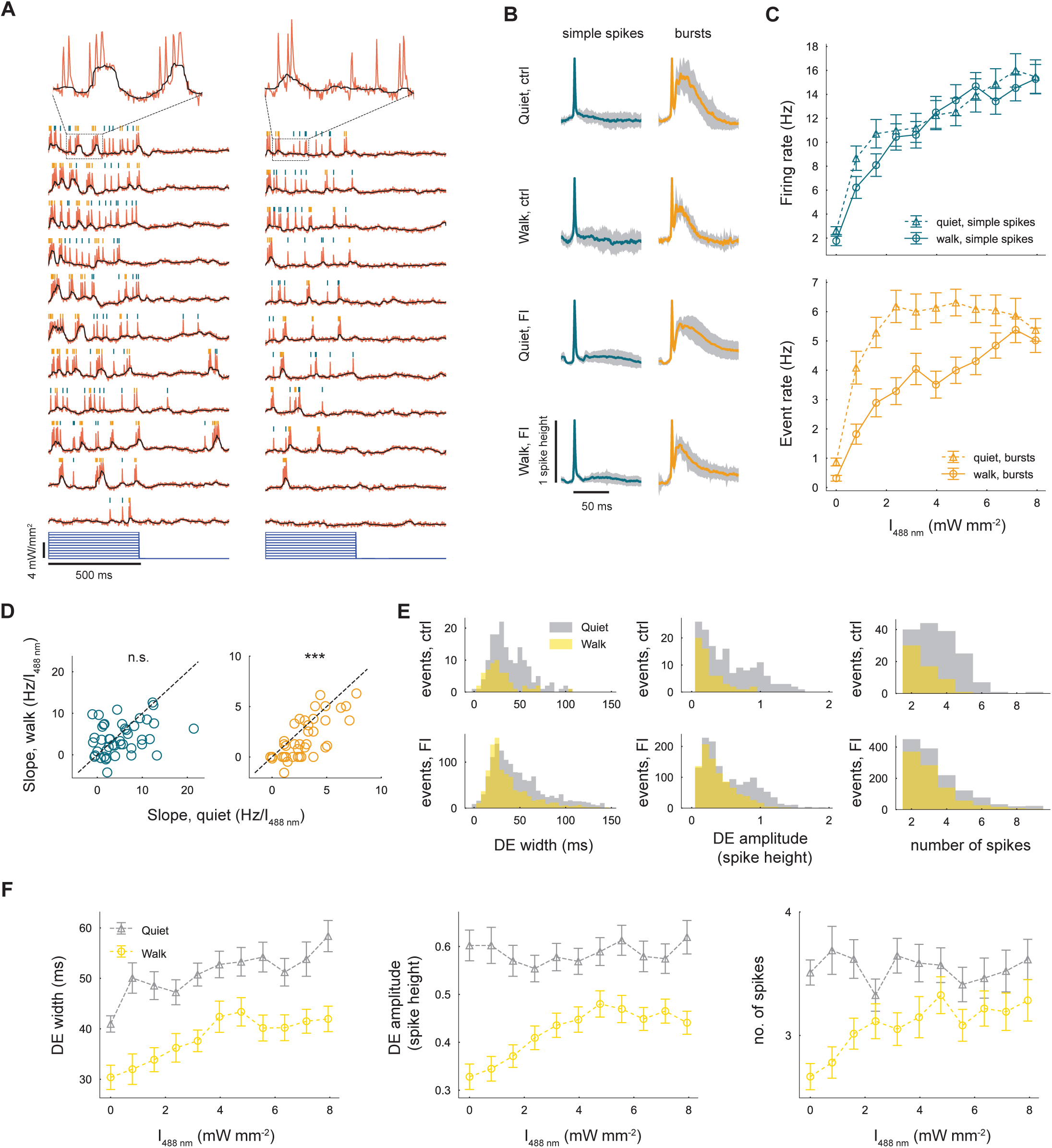
Burst-specific gain reduction in CA1 PCs during walking. (A) Optically recorded activity of a CA1 PC during the optogenetic F-I protocol, showing simple spikes (teal) and bursts (orange). Black traces represent subthreshold Vm. (B) Population-averaged spike waveforms for simple spikes (left) and bursts (right) under various experimental conditions. Shaded areas represent the s.e.m. (C) F-I curves for simple spikes (top) and bursts (bottom). (D) Scatter plots comparing slopes of F-I curves for simple spikes (left) and bursts (right) during quiet and walking. (E) Distribution of DE width (left), DE amplitude (middle), and the number of spikes per burst (right) in quiet (gray) and walking (gold) conditions, before (top) and during (bottom) the F-I protocol. (F) Effect of stimulation intensity on DE width (left), DE amplitude (middle), and the number of spikes per burst (right) in quiet and walking conditions during the F-I protocol (data are mean ± s.e.m.).

Plateau potentials have recently been implicated in behavioral-time-scale plasticity (BTSP) and are thought to play a crucial role in the formation of new place fields during locomotion^15,40^. Therefore, we quantified various aspects of the DE across behavioral states. We found that walking significantly reduced DE width (quiet ctrl., 41.1 ± 1.6 ms; walk ctrl., 30.4 ± 2.4 ms; p = 1.2×10^-4^; quiet FI, 52.1 ± 0.9 ms; walk ctrl., 39.8 ± 0.9 ms; p = 1.3×10^-23^, Mann-Whitney U test), amplitude (quiet ctrl., 0.60 ± 0.03 spike height; walk ctrl., 0.33 ± 0.03 spike height; p = 9.5×10^-7^; quiet FI, 0.58 ± 0.01 spike height; walk ctrl., 0.44 ± 0.01 spike height; p = 1.1×10^-18^, Mann-Whitney U test), and the number of spikes associated with each event (quiet ctrl., 3.5 ± 0.1 spikes; walk ctrl., 2.7 ± 0.1 spikes; p = 5.6×10^-6^; quiet FI, 3.5 ± 0.05 spikes; walk ctrl., 3.2 ± 0.05 spikes; p = 4.2×10^-7^, Mann-Whitney U test; Figures 5E and S6C). Additionally, during quiet, only the DE width exhibited a linear increase with simulation intensity, while its amplitude and the number of spikes per event remained unchanged. In contrast, during walking all three measures linearly increased with laser intensity until saturation around 5 mW/mm^2^ (Figure 5F). These findings suggest that walking reduces the sensitivity of all aspects of the burst-associated DE to expand the dynamic range of neural responses.

### A two-compartment model captures the burst-specific gain modulation

We next sought to investigate the potential mechanisms underlying the walking-induced gain reduction of bursts in PCs. Previous work demonstrated that gain modulation of spiking output requires coordinated increases in excitatory and inhibitory synaptic inputs^37^. However, this mechanism cannot account for the output-type-specific gain modulation we observed (Figures 5 and S7A). Burst firing in hippocampal CA1 PCs has been shown to depend on dendrite-soma integration^42^, and is closely associated with plateau potentials, which involve voltage-gated Ca^2+^ channels, NMDA receptors, and coincident inputs from EC3 and CA3^15,41^. These findings suggest that burst-specific gain modulation may arise from nonlinear dendritic mechanisms. Based on this and the previously established mechanisms of gain modulation^37^, we hypothesized that walking-induced reductions in burst gain could result from dendrite-specific E/I interactions. To our knowledge, no existing biophysical models incorporate all the components necessary to reliably generate plateau potentials^17,34,43^. Consequently, we limited our analysis to explaining the spike-related gain modulation observed in PCs using a minimalistic model. We employed a two-compartment model. The somatic compartment was modeled as a standard leaky integrate-and-fire (LIF) unit, incorporating spike-triggered adaptation and a refractory period of 3 ms^44^. The dendritic compartment, which received backpropagating action potentials and generated plateau potential-like depolarizations, was similarly modeled with spike-triggered adaptation. Synaptic inputs to both compartments were modeled as time-varying conductances, and the somatic compartment received additional current, simulating targeted optogenetic stimulation (Figure 6A).

**Figure 6.**
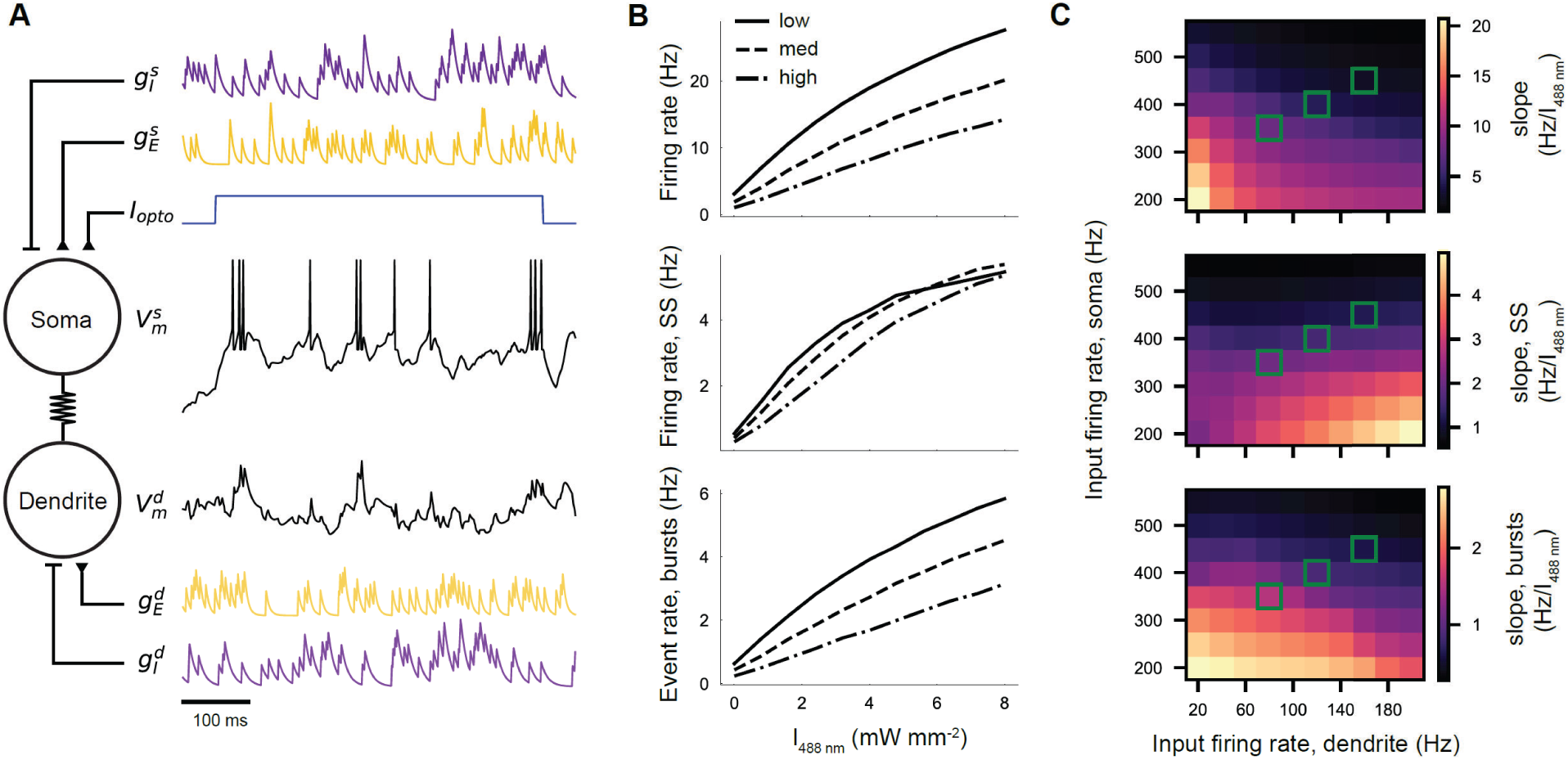
Burst-specific gain modulation in a two-compartment model neuron. (A) Schematic of a two-compartment model neuron receiving excitatory (yellow) and inhibitory (purple) synaptic inputs, with optogenetic current applied to the somatic compartment. The model includes excitatory and inhibitory synaptic inputs to both the somatic and dendritic compartments. The two black traces represent the somatic membrane potential (top) and dendritic membrane potential (bottom). (B) F-I curves for all spikes (top), simple spikes (middle), and bursts (bottom) under various input configurations. Low, medium, high indicate the strength of the excitatory and inhibitory synaptic input rates, corresponding to dendritic input frequencies of 80, 120, and 160 Hz, and somatic input frequencies of 350, 400, and 450 Hz, respectively. (C) Heatmaps illustrating the changes in the slopes of F-I curves for all spikes (top), simple spikes (middle), and bursts (bottom) as a function of dendritic and somatic input firing rates. Green squares highlight the synaptic input configurations displayed in (B).

Initially, we set the coupling between the soma and dendrite to zero and increased the firing rate of somatic synaptic excitatory and inhibitory inputs in a coordinated manner. This simulation reproduced the gain reduction observed in the F-I curve for all spikes (Figures 4C-4E), consistent with previous findings^37^. However, this gain modulation was observed primarily in simple spikes, as the single-compartment simulation essentially did not generate bursts (Figures. S7A and S7C). Our data suggest an increase in both somatic inhibition by PV INs and dendritic inhibition by SST cells during walking (Figure 2B). However, our model showed that shunting inhibition in both compartments resulted only in an additive shift of the F-I curves (Figure S7E), consistent with previous experimental findings and one-compartment model results^37^. Thus, we next simulated a coordinated increase in the excitatory and inhibitory input firing rates within both the dendritic and the somatic compartments. This adjustment qualitatively captured the experimental results, i.e. burst-specific gain reduction as observed during walking as well as minimal modulation of the simple spikes (Figure 6B). Notably, increasing dendritic input firing rates reduced burst gain while increasing simple spike gain, effectively transforming bursts into simple spikes (Figures 6C, S7B and S7D). However, further increasing the input firing rates in the somatic compartment counteracted the increase in the simple spike firing rate (Figures 6C, S7A and S7C). Overall, our simulation results suggest that a coordinated increase in the excitatory and inhibitory background firing rates in both the somatic and dendritic compartments could contribute to the specific gain modulation of bursts observed in our experimental data (Figure 5).

### A simple classifier precisely identifies the CA1 cell types based on their depolarized spike waveform

In this work, we generated a unique dataset with detailed membrane potential dynamics of four distinct cell types, each unambiguously identified through genetic labeling (Figures 1, S1 and S2). We thought to leverage these data to develop a computational method to infer the molecular identity of CA1 neurons based on their electrophysiological signatures.

Identifying cell types based on spike waveform is widely used in extracellular electrophysiology, though it is often unreliable and can typically separate only regular spiking from fast-spiking neurons^45^. Voltage imaging, however, offers additional insights by revealing subthreshold activities. Furthermore, the combination with optogenetic depolarization could expose additional intrinsic properties such as after-hyperpolarization. To capitalize on these signals, we analyzed the average spike waveform of all cells. We found these waveforms to be distinct across cell types and under various experimental conditions (Figure 7A). Therefore, we trained a one-dimensional convolutional neural network (1D-CNN)^46,47^ to classify cell types based on their average spike waveforms (Methods). The model performed optimally when trained on data from the walking and prolonged depolarization condition, achieving an average accuracy of 93% across the test dataset (Figure 7B). Analysis of the confusion matrix revealed that PV cells were identified with 100% accuracy, as expected from their distinct fast-spiking phenotype (Figure 7C). Strikingly, our methods provided high-accuracy identification also for the 3 regular spiking populations, reaching a 97% accuracy for PCs, 94% for SST cells, and 90% for VIP cells (Figure 7C). Notably, training the classifier using a minimal 40 ms spike window reached close to 90% accuracy (Figure S8), suggesting that the classification relies mostly on intrinsic biophysical cellular properties with little additional contribution from behavior-dependent activity patterns. This high-accuracy classification enables precise decoding of molecular cell type identities in wild-type (WT) animals, offering an invaluable tool for future studies.

**Figure 7.**
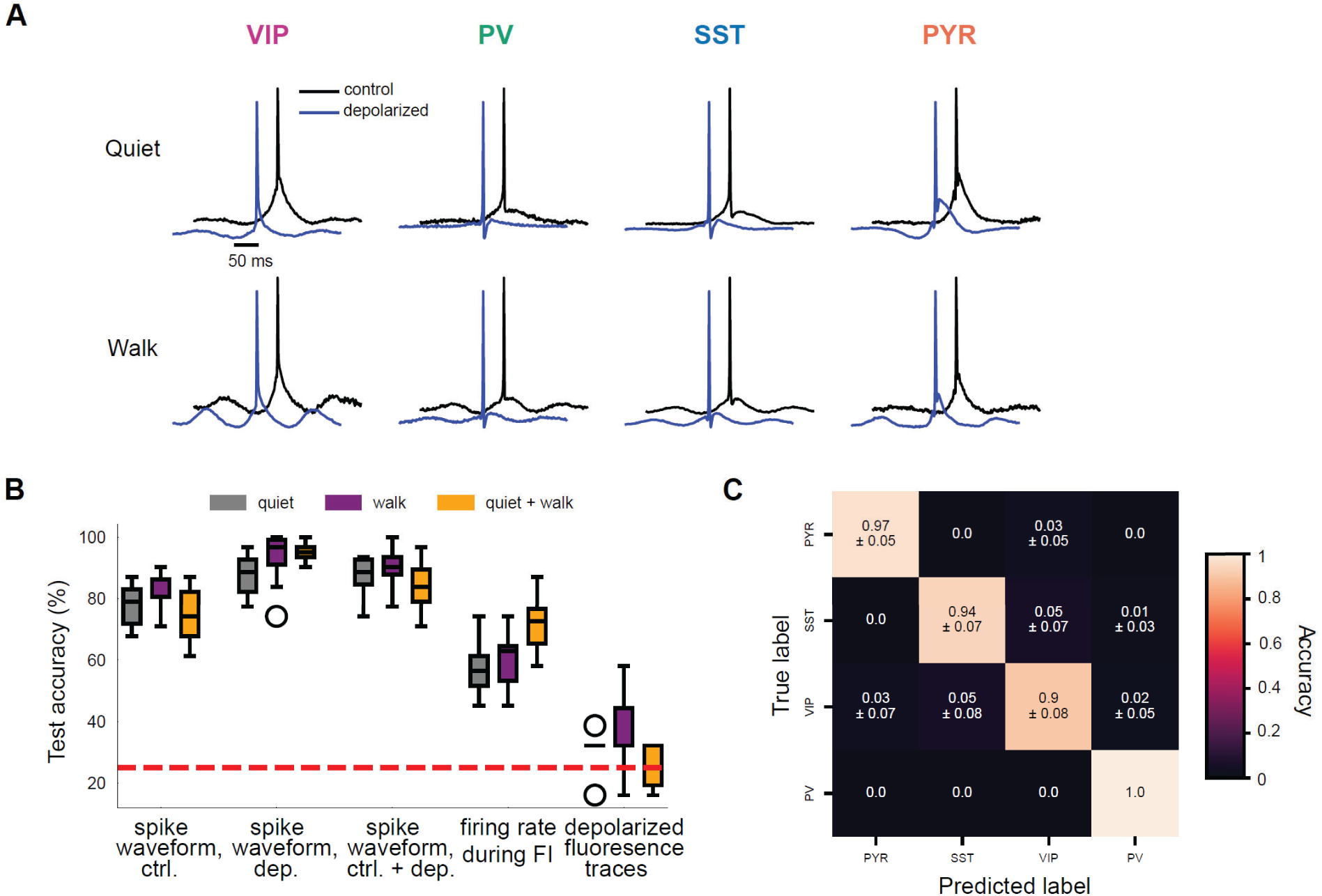
Decoding molecular identity with a 1-D CNN based on spike waveforms. (A) Grand average of spike waveforms for each cell type during quiet (top) and walking (bottom) states, with (blue) and without (black) optogenetic stimulation, illustrating both spiking and subthreshold characteristics. (B) Performance of the 1-D CNN trained on 70% of the experimental data for each cell type, evaluated across various data types, including spike waveforms and other forms of neural data, from the quiet state, the walking state, and a combination of both states. Results represent the mean and standard deviation of 10 random train-test splits. (C) Confusion matrix of the 1-D CNN trained with spike waveforms during walking with prolonged optogenetic stimulation, showing classification accuracy (mean ± s.d.) along the diagonal and misclassification probabilities in the off-diagonal elements.

## Discussion

In this study, we used cell-type specific all-optical electrophysiology in behaving mice and reported behavioral state-dependent modulation of local network firing patterns, subthreshold oscillatory activity, and neuronal gain in excitatory (E) and multiple inhibitory (I) cell types in CA1. By combining the recording of both local inhibitory cells and output excitatory PCs with optogenetic modulation at single-cell resolution and computational modeling, we showed how the dynamics of E and I synaptic inputs play a central role in shaping the hippocampal output.

Our results on the walking-induced firing rate decrease in PCs align with findings from intracellular recordings^23,35,48,49^. However, these results differ from those obtained using extracellular recordings and calcium imaging^50,51^. Several factors could explain this discrepancy, including differences in behavioral paradigms (we used controlled movement rather than voluntary movement, controlled the animal’s speed, and placed the animal in a dark environment with uniform tactile cues) or the recording techniques themselves. While this discrepancy highlights the influence of experimental conditions and recording techniques, our findings in three other CA1 IN types were consistent with the established local microcircuit connectivity^11^. The behavior-dependent firing pattern changes indicate that distinct input patterns are integrated across CA1 cell types. While we cannot infer changes in all input streams, we specifically uncovered how locomotion modulates theta-oscillating E or I inputs to each cell type by examining intracellular theta oscillation under prolonged depolarization and using a biophysical model. We found that PCs and VIP INs primarily receive inhibitory theta-modulated input, whereas excitatory theta-modulated input plays a more prominent role in PV and SST INs (Figure 3). As each cell type likely receives a combination of both E and I oscillatory inputs, which may or may not exhibit phase differences. we used simulations and demonstrated that the ratio between E and I inputs, as well as their relative phase difference, affects the intracellular theta modulation during prolonged depolarization (Figures 3D, 3E and S4).

The medial septum (MS), a well-known extrinsic theta pacemaker^10^, likely provides theta-oscillating inputs to CA1 through distinct projections: glutamatergic inputs to PV and SST INs^50^, GABAergic inputs to VIP INs^52^, and cholinergic inputs to PCs^53^. Theta oscillations can also originate intrinsically within the hippocampus^36^. Local CA1 PCs have been shown *in vitro* to drive theta activity in PV and SST INs^36^. Additionally, our findings demonstrated that VIP INs exhibit strong theta-modulated firing (Figure 3A), suggesting they provide local inhibitory theta input to PV and SST INs. For PCs, the main source of inhibitory theta inputs is likely local PV and SST INs (Figure 3A)^36^. Finally, our results and model imply that while EC3 and CA3 PCs contribute theta-modulated inputs to CA1 PCs^54^, these inputs are excitatory and thus play a relatively minor role in driving theta oscillations during controlled locomotion. In summary, we propose that PCs predominantly receive inhibitory theta inputs from local PV and SST INs and weak excitatory theta from CA3 and EC3, while VIP INs primarily receive inhibitory inputs from MS GABAergic neurons. PV and SST INs receive excitatory theta inputs from medial septal glutamatergic neurons along with inhibitory theta inputs from local VIP INs.

Our previous study^23^ showed that prolonged depolarization slightly enhanced the intracellular theta oscillations of SST INs, whereas here we observed a weak reduction. A key factor that could explain this discrepancy is the fact that in the previous study, we optogenetically stimulated all cells in the FOV, potentially inducing network-level effects, such as reduced feedback excitation onto SST INs due to PC inhibition. Here, in contrast, we minimized these effects by carefully applying single-cell stimulation. Additionally, our modeling results suggest that when theta-oscillating E/I conductances are nearly in phase with an I/E ratio around 3.5, the resulting currents are out of phase and tend to cancel each other out. This cancellation introduces high variability in the relative change in theta power, ranging from small enhancements to slight suppression, as the neuron becomes more sensitive to input noise (Figures S4E and S4F).

Based on the changes in firing rates across various cell types, one might intuitively attribute the reduction in PC firing during locomotion to the increased inhibition from PV and SST INs. However, our results from F-I curves and computational modeling (Figures 4c and 6) suggest a more nuanced mechanism. Specifically, we found that a coordinated increase in E/I inputs to PCs reduces spontaneous firing and gain. This effect on gain modulation could not be explained purely by inhibition but instead reflects an intricate balance between E and I inputs. Such insights are uniquely enabled by our method of measuring F-I curves across different behavioral states. By analyzing the F-I relationships in identified cell types during both quiet and locomotion, we uncovered three distinct modes of gain control: divisive in PCs, subtractive in VIP cells, and saturating in PV and SST cells. These modes have been reported in previous slice studies and shown to correspond to specific changes in E/I background input^37^. While changes in background E and I inputs are central to gain modulation, they are not the sole contributors. Network-level mechanisms, such as increased feedback inhibition, may also play a role in reducing gain^55^. Our results may also be partially explained by cholinergic input from the medial septum to both CA1 PCs and SST INs whose heightened activity during locomotion potentially depolarizes PCs while enhancing distal dendritic inhibition via SST INs^29^.

Furthermore, we observed that the gain reduction in PCs during locomotion is specific to bursting output. Dendritic mechanisms, such as the interaction of dendritic depolarization with back-propagating action potentials, have been implicated in gain modulation^42^. However, the dendritic mechanism alone modulates the gain of simple spikes and bursts simultaneously but in opposite directions, as one output is transformed into the other^42^ (Figures S7B and S7D). Our model results show that burst-specific gain reduction can be explained by a coordinated increase in E/I noisy inputs when applied to both somatic and dendritic compartments (Figure 6). Specifically, increased E/I noisy input to the dendritic compartment reduces the gain of bursts while increasing the gain of simple spikes, as bursts are transformed into simple spikes (Figure S7C). This gain increase of simple spikes in the dendrite is offset by a corresponding gain decrease in the somatic compartment (Figure S7D). Based on our modeling and experimental results, we propose that locomotion induces simultaneous increases in excitatory inputs from EC3 and CA3, along with inhibitory inputs from SST and PV INs, resulting in reduced burst gain in PCs. While increased SST and PV INs activity was directly observed in our data (Figure 2), we lack direct evidence for changes in EC3 and CA3 inputs to PCs. However, local cortical circuits generally generate proportional inhibition in response to excitation^1–3^, and hippocampal slice studies have demonstrated that heightened feedforward excitation from EC3 and CA3 increases the activity of SST and PV INs, respectively^32^. Additionally, silencing both SST and PV INs significantly increases burst firing, whereas silencing them individually has shown conflicting results^32,56^. This highlights the necessity of both compartments in modulating burst activity. Therefore, a simultaneous increase in excitatory and inhibitory synaptic inputs in both somatic and dendritic compartments serves as a plausible mechanism for burst-specific gain reduction, as indicated by our two-compartment model.

A closer examination of burst-associated depolarization revealed that walking induces shorter, smaller depolarization with fewer burst spikes (Figures 7B and 7E), many of which potentially correspond to calcium plateau potentials generated in the dendrites that are critical for plasticity mechanisms underlying place field formation^15,40^. Based on our data and model, we interpret the walking state as a default exploration mode in an environment devoid of behaviorally salient cues. In this state, spatially uniform excitation from CA3 and EC3, coupled with uniform inhibition from SST and PV interneurons, reduces the gain of burst firing and the generation of plateau potentials in PCs. In contrast, sensory cues likely induce non-uniform excitatory inputs from EC3 to individual CA1 PCs, which, when coinciding with CA3 inputs, result in plateau potentials at specific locations^15,57^. Sufficiently large and prolonged plateau potentials can trigger BTSP and promote the formation of distinct place fields^15,25,40^. Consequently, CA1 PCs transition from a default E/I balanced mode to an excitation-dominant mode within place fields^13^. Furthermore, the properties of burst-associated depolarizations linearly change across a wider range of inputs during walking, in contrast to the limited changes in duration observed during quiet. This effect is possibly mediated by enhanced local inhibition^17,48^, enabling finer control over plasticity induction. The reduced sensitivity of DE responses, combined with the burst-specific gain reduction during locomotion, might therefore enhance in-field selectivity and promote more precise spatial coding when the mouse encounters rich sensory cues in a cognitively engaging environment^15,17,31^. Additionally, this mechanism may allow CA1 PCs, as a network to achieve a sparse representation of the environment, enhancing memory storage capacity^30^.

Lastly, our cell type classifier, trained from data combining voltage imaging and optogenetic stimulation, provides a powerful posthoc method to accurately identify four distinct CA1 cell types. This approach enables simultaneous monitoring of large populations of neurons during behavior, offering detailed insights into the interactions between multiple cell types. Moving forward, combining all-optical electrophysiology with posthoc cell type identification presents exciting opportunities to study neuronal activity during learning and memory formation. Applying these tools in more cognitively demanding tasks could illuminate how distinct cell types contribute to circuit E/I balance, information processing, spatial representation, behavioral or contextual adaptation, and *in vivo* plasticity.

## Resource availability

### Lead contact

Further information and requests for resources should be directed to and will be fulfilled by the lead contact, Yoav Adam

## Material availability

This study did not generate new unique reagents.

## Data and code availability

All data reported in this paper will be shared by the lead contact upon request.

All original code will be deposited at GitHub.

Any additional information required to reanalyze the data reported in this paper is available from the lead contact upon request.

## Acknowledgments

We thank Haim Somplonisky and Jonathan Kadmon for useful discussions at the early stages of this project. We thank Gili Kupferman and members of the Adam lab for useful discussions. We thank Maya Groysman and the ELSC vector core facility (EVCF) team for virus production, and Itamar Frachtenberg and the ELSC Fab Lab team for help with the design and fabrication of the behavioral setup. This work was supported by the European Research Council (ERC) starting grant #948716 and the Israel Science Foundation (ISF) grant #2940/24 to YA. YA is the Sachs Family Lecturer in Brain Sciences.

## Author contributions

QY performed all surgeries, *in vivo* imaging experiments, and data analysis, with help from YA and RK. SBS performed all the molecular biology and histology. QY developed the computational models and simulations with advice and supervision from ML. YA provided funding and supervised all aspects of the project.

## Supplemental information

Document S1. Figures S1–S8

## Methods

### Animals

Male and female mice aged 12–18 weeks, including 3 SST-Cre (Jax #013044), 3 VIP-Cre (Jax #010908), 3 PV-Cre (Jax #017320), and 3 CKII-Cre (Jax #005359) mice, were used. All mice were heterozygous and maintained on C57BL/6 background. All animal experiments were conducted in accordance with guidelines approved by the Institutional Animal Care and Use Committee (IACUC) of the Hebrew University of Jerusalem.

### Surgery

The surgical procedures followed protocols similar to those described previously^23^. Mice aged 12–18 weeks were anesthetized with 2% isoflurane for induction and maintained at approximately 1% isoflurane during surgery. Mice were positioned in a stereotaxic frame, and ophthalmic ointment was applied to prevent corneal drying. A regulated heating pad maintained the body temperature at 37°C. The surgical site was exposed by retracting the scalp, after which the skull surface was rinsed and dried. A 3-mm diameter craniotomy was performed over the right dorsal CA1 region (ML: 1.8 mm, AP: 2.0 mm) using a 3-mm biopsy punch (Miltex). The dura mater was carefully removed, and the underlying cortex was aspirated under continuous saline irrigation. The external capsule was then exposed, and a small central portion was delicately removed to expose the CA1 area.

Using a microsyringe attached to a UMP3 microinjection pump (World Precision Instruments), specific viral constructs were delivered at a rate of 1 nl/s from −0.2 to 0 mm relative to the exposed CA1 surface, totaling 80 nl per injection depth. For SST-Cre and VIP-Cre mice, hSyn-DIO-somArchon1-P2A-somCheRiff-HA virus (final titer 4.5 × 10^12^ GC/ml) was injected. CKII-Cre mice received a mixture of hSyn-fDIO- SomArchon1-P2A-somCheRiff-shortPA virus (final titer 4 × 10^12^ GC/ml) and EF1a-DIO-FlpO (final titer 1.72 × 10^10^ GC/ml) for sparse expression. For PV-Cre mice, hSyn-FLEx -SomQuasAr6b-P2A-somCheRiff-HA (final titer 2.4 × 10^12^ GC/ml) was used to monitor fast spikes of PV INs. All viruses were produced at the ELSC vector core facility (EVCF).

Following injection, a custom-made cannula was prepared using a 2.0-mm segment of 3-mm diameter stainless steel tube (MicroGroup), capped with a no. 1 round cover glass using UV curable glue (NOA81, Norland). The cannula was implanted and secured to the skull with 3M Vetbond tissue adhesive. After the cannula was cured, a custom-made titanium head plate was glued around the cannula, and any exposed skull was covered with dental cement (C&B Superbond, Sun Medical). Animals were returned to their cages for recovery and treated with meloxicam (5 mg kg^-1^) for 2 days.

### Behavior

During imaging sessions, the animals were head-fixed on a 3D-printed wheel and their behavior was controlled by an electric motor. Each field of view was imaged during motor-off quiet periods for 35 seconds, and then while walking at a constant speed of 10 cm/s for 35 seconds. Prior to imaging, and at least 2 weeks after the surgery, the animals were habituated to both head restraint and controlled walking. The habituation process involved training the animal over at least three sessions, each lasting 15 minutes. During these training sessions, animals were initially allowed to relax on the wheel. Following this acclimation period, they were subjected to a training protocol consisting of 1-minute walking intervals followed by 2-minute rest intervals. The walking speed during training was gradually adjusted to range from 5 to 10 cm/s to ensure proper adaptation to the experimental conditions. The last training session was conducted under the microscope with an objective above the cranial window.

### Optical setup

Imaging was conducted on an upright Bergamo microscope body equipped with a scanning two-photon module (not used in this study) and a widefield epifluorescence module (Thorlabs). The microscope was also equipped with a custom holography module designed and built by Thorlabs Imaging Systems Inc. according to our specifications. The module included a 120 mW CW 488 nm laser (Coherent) and 1 W 639 nm CW laser (CNI). Both lasers were fiber-coupled and were patterned using a spatial light modulator (SLM, EXULUS-HD2, Thorlabs). Each beam was expanded, collimated, passed through a ¼ wave plate, and targeted to half of the SLM. The two wavelengths were then split and recombined using a pair of dichroic mirrors and relayed to the objective back aperture (Figure 1B). The zero’s order was blocked in an intermediate image plane using a glass with a deposited gold point. Mice were imaged using a 16X 0.8 NA 3-mm working distance objective (Nikon, N16XLWD-PF) and a custom 252 mm tube lens (effective magnification 20.1X), except for PV-Cre mice which were imaged with a 25X 1.0 NA 4-mm working distance objective (Olympus, XLPLN25XSVMP2) and the same tube lens (effective magnification 35X). The red laser intensity was 24 W mm^-2^ at the sample. The blue laser was attenuated using a MEMS-baed electronic attenuator (V450A, Thorlabs), and cells were stimulated at 0.8 to 8 mW/mm^-2^ (Figure 1D and Figure 4). Fluorescence was collected on a scientific CMOS camera (Hamamatsu, ORCA-Fusion) at a sampling rate of 1000 frames/sec.

### Simultaneous voltage imaging and optogenetic stimulation in behaving animals

We implemented a 71-second protocol (Figure 1D, top) starting with a 6-second resting baseline, followed by a 10-step F-I protocol with optogenetic stimuli increasing from 0.8 to 8 mW/mm², each applied for 500 ms with 500 ms intervals. Eight seconds after the F-I protocol, an 8-second constant optogenetic stimulus at 4 mW/mm² was applied. Four seconds after this stimulus ended, the motor was activated, and 5 seconds after baseline recording, the same F-I, and prolonged optogenetic stimuli were repeated during controlled walking. In each field of view (FOV), we recorded up to 5 PCs or 3 SST INs simultaneously. Due to photobleaching limitations, we applied the protocol up to 3 times per FOV, stimulating only one cell at a time. For VIP and PV INs, only one cell was recorded and stimulated once per FOV. An imaging session lasted up to 2 h and then animals were returned to their home cages.

### Immunohistochemistry

Animals were deeply anesthetized with isoflurane and then perfused transcardially with ice-cold PBS (pH 7.4) followed by 4% paraformaldehyde in PBS. After overnight post-fixation in 4% paraformaldehyde, brains were moved to PBS. Coronal sections {50 µm) were cut on a Vibratome (VT1000S, Leica), collected in PBS, and stored at 4 °C for further use. Primary antibodies used were rat anti-somatostatin (1:50, Merck MAB354), and rabbit anti-parvalbumin (1:100, Abcam ab11427). Secondary antiboeis used were Alexa Fluor 594 donkey anti-rat IgG (1:500, Abcam), and Alexa Fluor 647 goat anti-rabbit IgG (1:500, Abcam). Following 1 h in PBS containing 5% goat serum and 0.1% Tritin X-100, sections were incubated with primary antibodies for 36–48 h. Following washes in PBS, slices were incubated in secondary antibodies for 1 h, and then mounted with DAPI containing mounting medium (Vectashield). Confocal images were acquired with Leica Stellaris 5 microscope equipped with a white light super-continuum laser.

### Image segmentation and signal extraction

The signal extraction pipeline was modified based on the VolPy algorithm integrated into the CaImAn package (https://github.com/flatironinstitute/CaImAn)^58^. Briefly, the movies were first motion-corrected using the NoRMCorre algorithm^59^. We used a kernel size of 3 for high-pass spatial filtering and rigid motion correction. Movies were segmented based on the binary masks used for patterned illumination during the experiment. Each segment of the movie was corrected for photobleaching through high-pass filtering. An initial fluorescence trace was extracted by averaging the pixels within the region of interest (ROI) indicated by the binary mask. Background noise was removed through singular value decomposition (SVD) and ridge regression. The denoised trace was then passed through an adaptive thresholding algorithm to extract prominent spikes. The waveforms of these extracted spikes were averaged to generate a template, which was then used in a whitened-matched filter to detect the spikes again. The resulting spike train was convolved with the template to generate a spiking trace. Next, a weighted spatial footprint (SPF) was generated by ridge-regressing the spiking trace on the high-pass filtered movie. This SPF was then applied to the segmented movie again to generate a trace with a better signal-to-noise ratio (SNR). The process of spike detection and SPF refinement was repeated once more on the improved trace, and the resulting SPF was used to extract the optimized fluorescence trace.

### Data analysis

#### Spike detection and subthreshold Vm extraction

Due to the potential reduction of SNR, we divided the fluorescence trace into two halves and conducted spike detection separately for each half. For VIP, PV, and SST INs, spikes were extracted from the fluorescence trace using the built-in spike detection algorithm from Volpy as described above.

For PCs, which often exhibit complex spikes with different waveform shapes compared to simple spikes, we used a simple thresholding algorithm to extract the spikes in the final round. A custom graphical user interface was developed to manually adjust the threshold and inspect the quality of the resulting spikes. We excluded slow spikes that typically occurred at the end of a burst, as their waveforms were distinct from simple spikes, characterized by a wider spike width and smaller amplitude in the high-pass filtered trace. To achieve this, we applied a whitened matched filter based on the average waveform of simple spikes. Spikes with amplitudes in the filtered trace lower than 70% of the simple spikes’ average amplitude were excluded. This step did not qualitatively change the F-I results.

To extract the subthreshold Vm from the optimized fluorescence trace, we typically removed 3 data points before and after each spike peak, linearly interpolated the missing data, and then applied a 5 ms moving average filter for smoothing. For detecting burst-associated DEs in PCs, we median filtered the fluorescence trace (25 ms) instead, as this method better preserves the shape of the DEs.

#### Subthreshold Vm power spectra and mean theta amplitude

To compare power spectra and theta amplitude across conditions, we normalized the subthreshold Vm by the average spike amplitude calculated from the fluorescence trace for each condition separately. The spike amplitude of each spike was measured by the difference between the peak spike fluorescence and the lowest fluorescence value within 5 ms before the peak. The power spectra were calculated as the real part of the product of the Fourier transform of the normalized trace and its complex conjugate. The mean theta amplitude was calculated by taking the average of the amplitude obtained from the Hilbert transform of the subthreshold Vm, after band-pass filtering it between 6 and 9 Hz.

#### F-I Curve Fitting and slope calculation

To model the relationship between laser intensity (I) and neuronal firing rate (F), we used a sigmoidal function that captures the saturating nature of neuronal responses to increasing stimulus intensity and accounts for the variable spike-evoking laser intensity threshold observed in the F-I curves:

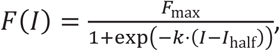

F_max_ represents the maximum firing rate, which is the asymptotic value the firing rate approaches as laser intensity increases. The parameter *k* determines the steepness of the curve. The I_half_ parameter is the laser intensity at which the firing rate is half of its maximum value. Cells with goodness of fit greater than 0.1 in both quiet and walking conditions were included in the grand averaged F-I curves and the calculation of the F-I slope.

To calculate the F-I slope for all spikes, we identified the most linear portion of the fitted sigmoidal curve. For most cells, the left boundary of this linear region was set at the point where the laser intensity was 0, while the right boundary was defined as the laser intensity that evoked 80% of the maximum firing rate. However, in cases where the F-I curve exhibited a high threshold (see examples in Figure S6A), we adjusted the left boundary to the point where the laser intensity produced more than 20% of the saturation firing rate. Similarly, if the F-I curve did not reach 80% of the saturation firing rate, we set the right boundary at the highest laser intensity. We obtained the slope by performing linear regression on this portion of the F-I curve between the left and right boundaries.

To calculate the F-I slope for simple spikes and bursts in PCs, we observed that most output-specific F-I curves did not exhibit a sigmoidal shape but appeared approximately linear across the first three data points. Therefore, we calculated the slope of the F-I curves by performing linear regression on these initial three data points.

#### Burst detection

We adapted our burst detection method from a recent study^34^. Putative bursts were identified using an ISI threshold of 14 ms. For each detected burst, the baseline was defined as the minimum value among five data points of the subthreshold Vm (obtained via median filtering with a 25 ms window) preceding the first spike within the burst. The local spike height was calculated as the difference between the raw fluorescence of the first spike and the baseline. A threshold of 0.15 spike height above the baseline was applied, and we assessed whether the subthreshold Vm crossed this threshold before the final spike in the burst. If the threshold was crossed, the event was classified as a burst; otherwise, all spikes within the event were classified as simple spikes.

We observed that the putative bursts associated with pronounced DE often exhibited ISIs exceeding 14 ms between the final spikes (Figure S6A). To address this, we applied a merging step to include spikes within the depolarization window before the detection step. Specifically, we applied a higher threshold of 0.4 spike height above the baseline to define the DE window, with its boundaries set by the left and right crossing points of this threshold. All spikes within the defined DE window were merged into the burst.

To quantify DE duration, we used the 0.15 threshold to delineate the event window and calculated its duration as the difference between the right and left crossing points. DE amplitude was measured as the maximum value within the window minus the baseline and was subsequently normalized to the local spike height.

### Cell type classifier

We implemented a 1D CNN using PyTorch for cell-type classification tasks similar to previously published methods^46^. The network architecture included four convolutional layers with 256, 128, 64, and 32 filters respectively, each followed by batch normalization and ReLU activation. Max-pooling layers with kernel size 3, stride 2, and padding 1 were applied after the second, third, and fourth convolutional layers. A dropout layer (p=0.5) was included for regularization. The output was flattened and passed through a fully connected layer, followed by a softmax activation to produce class probabilities.

The dataset included average spike waveforms in the control condition, average spike waveforms when the cells were depolarized, firing rate during the FI protocol, and raw fluorescence traces from four cell types: SST (n=50), PV (n=21), VIP (n=28), and PYR (n=48), all data were z-score normalized. We randomly split the dataset 10 times, each dataset was split 70% for training and 30% for testing. To address the imbalanced dataset, we used weighted cross-entropy loss. The model was trained with a batch size of 16 and a learning rate of 0.01. We reported the mean accuracy and standard deviations of the 10 random train-test splits. Additionally, we evaluated performance using the leave-one-out method, where each instance in the dataset was used once as a test sample while the remaining instances formed the training set. All computations were performed on a Quadro RTX 8000 GPU with 48GB RAM.

### Models and simulations

#### Single-compartment CA1 pyramidal neuron model with theta oscillation input

We implemented a single-compartment model (adapted from the original Hodgkin-Huxley model) to simulate the CA1 neuron dynamics^60^. The current balance equation for this model is:

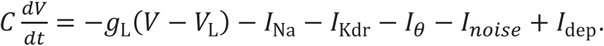

The neuron model parameters include a membrane capacitance C of 1 µF/cm^2^, a constant leak conductance *g*_L_ of 0.05*mS*/*cm*^2^, and a leak reversal potential *V_L_* of −70 *mV*. The active ionic currents include sodium and potassium channels, with maximal conductances of 35 mS/cm^2^ and 6 *mS*/*cm*^2^, respectively.

The theta oscillating synaptic current *I_θ_* is given by 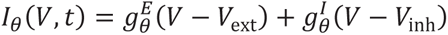, where the excitatory reversal potential *V*_ext_ = 0 *mV*, the inhibitory reversal potential *V*_inh_ = −80 *mV*. For figure 3d, 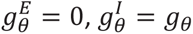. For figure 3e, 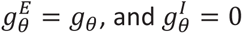. The theta oscillating conductance *g_θ_* is described by 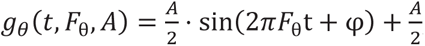, where t is the time, *F*_θ_ = 8 *Hz* is the theta frequency, and *A* is the peak amplitude. For figure 3d and 3e, *A* is set to 0.4 *mS*/*c*m^2^ and 0.1 *mS*/*c*m^2^, respectively. For Figure S4, the phase φ of E/I currents was varied to control the phase difference between them. The I/E ratio was set between 0 and 10 by fixing the peak E theta amplitude to 0.05 *mS*/*c*m^2^ and varying peak I theta amplitude.

The noisy current 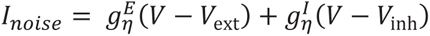 results from time-varying conductances 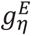 and 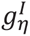, which are derived from the convolution of a synaptic conductance kernel with independent E/I Poisson spike trains, each with a firing rate of 200 Hz. The conductance has an instantaneous rise for both excitatory and inhibitory inputs, followed by a falling phase characterized by a 5 ms time constant for excitatory inputs and a 10 ms time constant for inhibitory inputs. The peak unitary conductances were set to 0.08 *nS* for both E and I noisy inputs.

The depolarizing current *I*_dep_ is set to 0 in the control condition, and *I*_dep_ = 0.2 *nA* in the depolarization condition to simulate the effect of prolonged optogenetic stimulation.

The simulations were performed using the fourth-order Runge-Kutta method, with a step size of 0.1 ms and total duration of 2 seconds implemented in Python.

We calculated the theta power *P_θ_* as the sum of PSD between 7-9 Hz and reported the normalized difference in theta power as 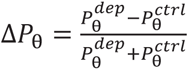.

#### Two-compartment model for modelling output-specific gain modulation

We implemented a two-compartment model to simulate the behavior of a CA1 pyramidal neuron^44^.

The somatic compartment, described by the membrane potential 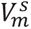, follows a leaky integrate-and-fire (LIF) model with spike-triggered adaptation. The membrane potential 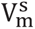 evolves according to the following equation:

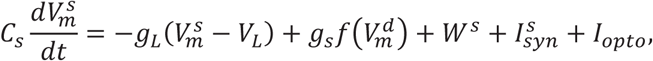

where C_s_ = 20 pF represents the somatic capacitance, and g_L_ = 1 nS is the leak conductance. The resting membrane potential is V_L_ = −65 mV. When 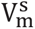 crosses the action potential threshold −54 mV, the membrane potential is reset to −60 mV, and a spike is generated, followed by a refractory period of 3 ms.

The coupling between the dendritic and somatic compartments is mediated by the term 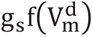, where g_s_= 1300 pA represents the coupling strength, and 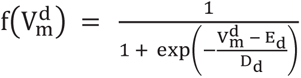 is a sigmoid function that captures the nonlinear influence of the dendritic membrane potential on the soma. Here, E_d_ = −38 mV is the midpoint, and D_d_ = 6 mV determines the steepness of the function.

Spike-triggered adaptation in the soma is modeled by the adaptation current W^s^, which evolves according to:

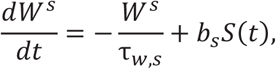

where the spike train *S*(*t*) = Σ*_i_*(*t* - *t_i_*), τ_W,s_ = 100 ms is the adaptation time constant, and *b_s_* = −10 pA controls the strength of the adaptation following each spike in the somatic spike train *t_i_*.

The synaptic current 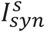 in the somatic compartment is determined by the excitatory and inhibitory synaptic conductances, 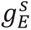 and 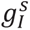, respectively^37^. These conductances are generated by presynaptic spike trains, with each presynaptic spike contributing to the conductance via a difference of exponential function. The kernel definitions and parameterizations are the same as above. The peak unitary conductances were set to 0.2 *nS* for excitatory and 0.8 *nS* for inhibitory inputs. The resulting synaptic current is given by 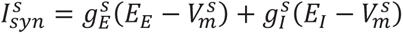, with reversal potential E_E_= 0 mV and E_I_= −80 mV. These synaptic inputs are driven by presynaptic spike trains generated by independent Poisson processes at specified rates. For Figure 6B and Figure S8A, to balance the excitatory and inhibitory synaptic currents, the inhibitory and excitatory presynaptic inputs to the somatic compartment have the same firing rates^37^: 350 Hz for the low condition, 400 Hz for the medium condition, and 450 Hz for the high condition.

The optogenetic intensity of the laser Γ*_opto_* is analogous to our experimental F-I protocol, it consists of 10 steps, each lasting 500 ms with 500 ms off intervals between them. Γ*_opto_* increases linearly from 0.8 to 8 mW/mm² across the steps, beginning after a 1-second delay with no stimulation. Γ*_opto_* was converted to optogenetic current I_opto_ by 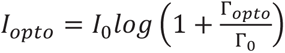, where *I*_0_ = 20 pA, and Γ_0_ = 0.8 mW/mm² are the parameters^61^.

The dendritic compartment, characterized by its membrane potential 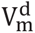, captures the generation of plateau potentials (or calcium spikes). The dendritic dynamics are governed by:

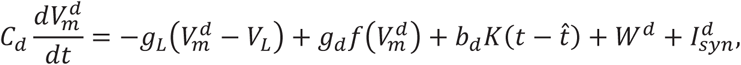

where C_d_ = 10 pF is the dendritic capacitance, and g_d_ = 14 pA modulates the strength of the dendritic nonlinear response. The term b_d_ = 38 pA influences the backpropagating action potentials (bAPs) following a somatic spike, modeled by the boxcar kernel 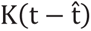.

The dendritic adaptation current W^d^ evolves as:

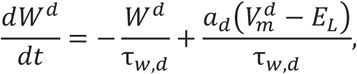

where τ_W,d_ = 30 ms is the adaptation time constant, and a_d_ = −13 nS controls the adaptation’s response to deviations in the dendritic membrane potential from the resting state. The dendritic synaptic inputs are given by 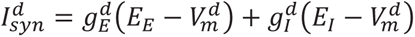. The generation of excitatory and inhibitory synaptic conductances follow the same description as those in the somatic compartment. For figure 6b, the inhibitory and excitatory presynaptic inputs to the dendritic compartment have the same firing rates as well: 80 Hz for the low condition, 120 Hz for the medium condition, and 160 Hz for the high condition.

The model results are based on 5000 simulations, each conducted using Euler’s method with a time step of 1 ms. We separated the spikes into simple spikes and bursts based on an ISI threshold of 14 ms.

### Statistical tests

Throughout the study, for paired comparisons, we first assessed whether the difference between two groups was normally distributed using the Shapiro-Wilk test. If the difference was normally distributed, we applied a paired t-test; otherwise, we used a Wilcoxon signed-rank test. For non-paired comparisons, we used Mann-Whitney U tests.

---

**Figure S1.**
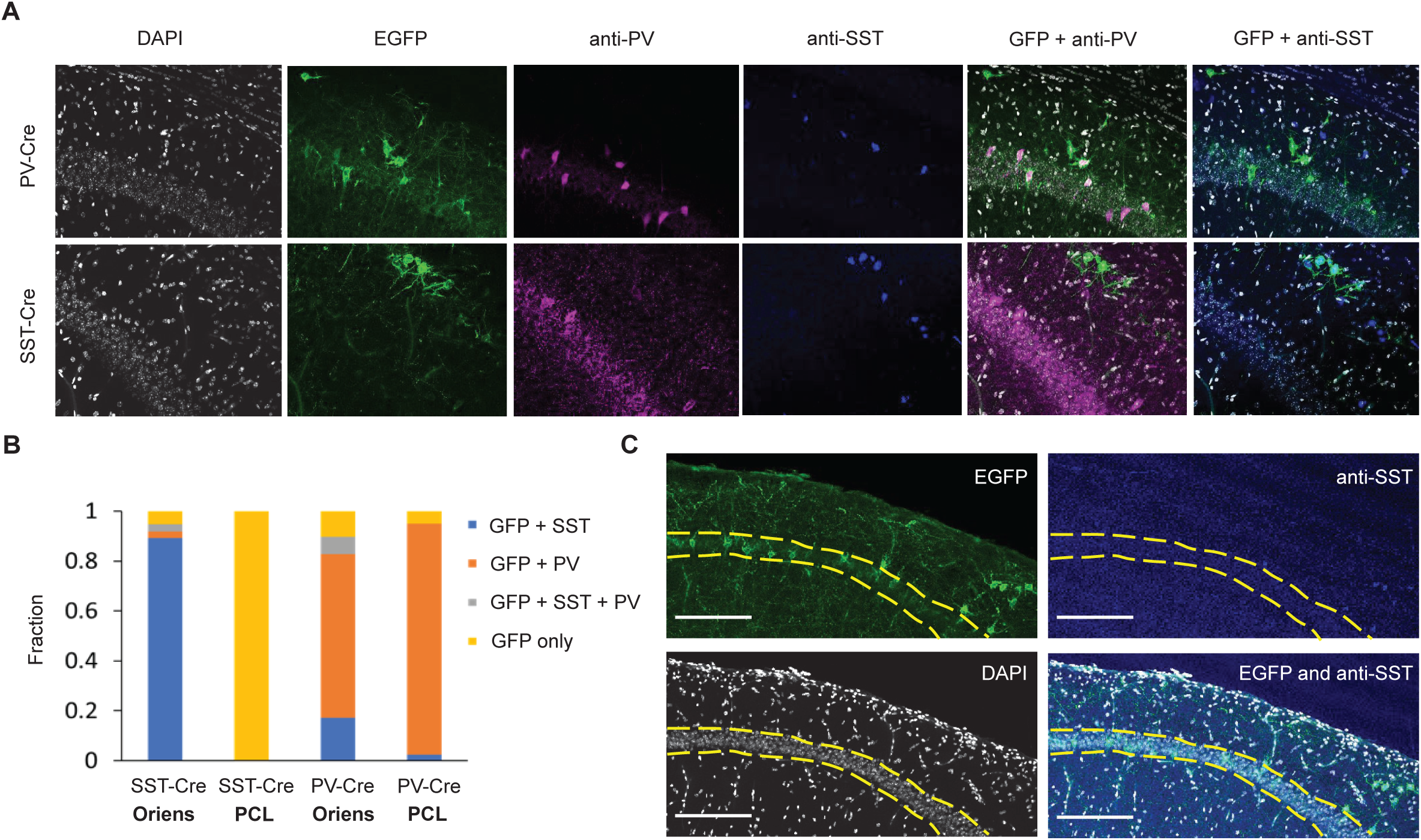
Co-immunostaining demonstrates anatomical distinctions between SST and PV INs, related to Figure 1. (A) Confocal images from PV-Cre and SST-Cre mice showing distinct populations of PV-positive and SST-positive cells, respectively, with co-expression of the Cre-dependent EGFP marker. PV-positive cells are observed in both the oriens layer and the pyramidal layer of the hippocampus, while SST-positive cells are primarily in the oriens layer. Scale bars, 90 μM. (B) Quantification of co-localization of EGFP with PV, SST, or both markers across different layers of the hippocampus in SST-Cre (N = 2) and PV-Cre (N=2) mice. The fraction of GFP-positive cells that co-express SST, PV, both, or neither marker is presented for the oriens layer (OL) and pyramidal layer (PCL) in each mouse line. Left bars represent SST-Cre mice (n = 37 cells in OL, n = 10 cells in PCL), and right bars represent PV-Cre mice (n = 29 cells in OL, n = 40 cells in PCL). (C) In some SST-Cre mice, leakiness is occasionally apparent as EGFP-positive cells in pyramidal layer, but leaky cells do not positively stain for SST or PV. Note that we did not image PCL neurons in SST-Cre mice. Dashed lines mark pyramidal layer borders. Scale bars, 180 μM.

**Figure S2.**
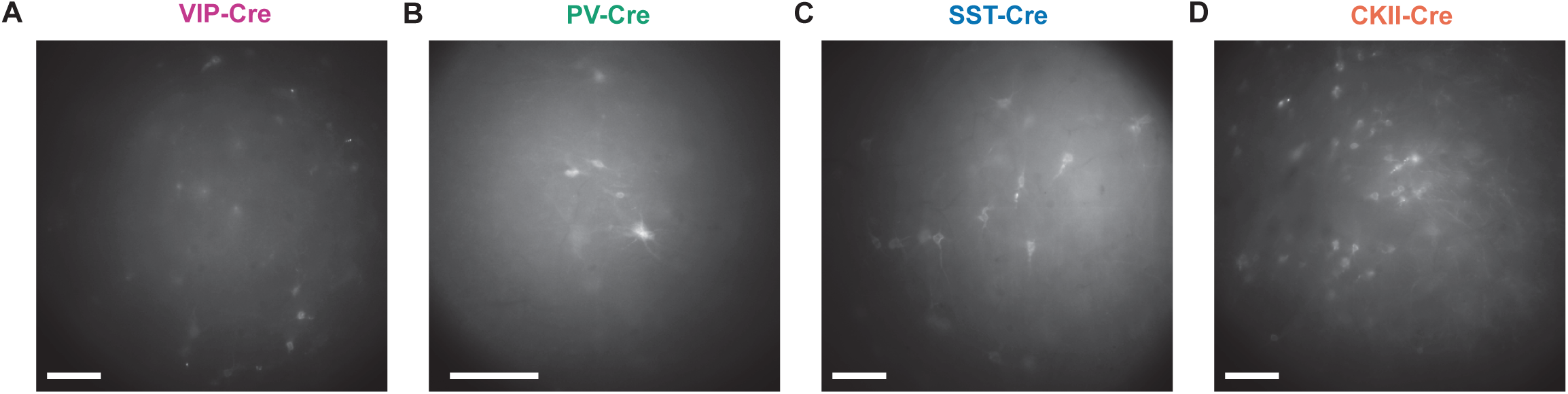
Sparse expression pattern of GEVI in vivo across 4 cell types, related to Figure 1. (A) Epifluoresence image with wide-field blue illumination of GFP under a 16x objective, showing expression pattern of the optopatch construct. Scale bar, 100 µm. (B) Same as (a), but under a 25X objective. (C-D) Same as (A), but for SST-Cre mice and CKII-Cre mice, respectively.

**Figure S3.**
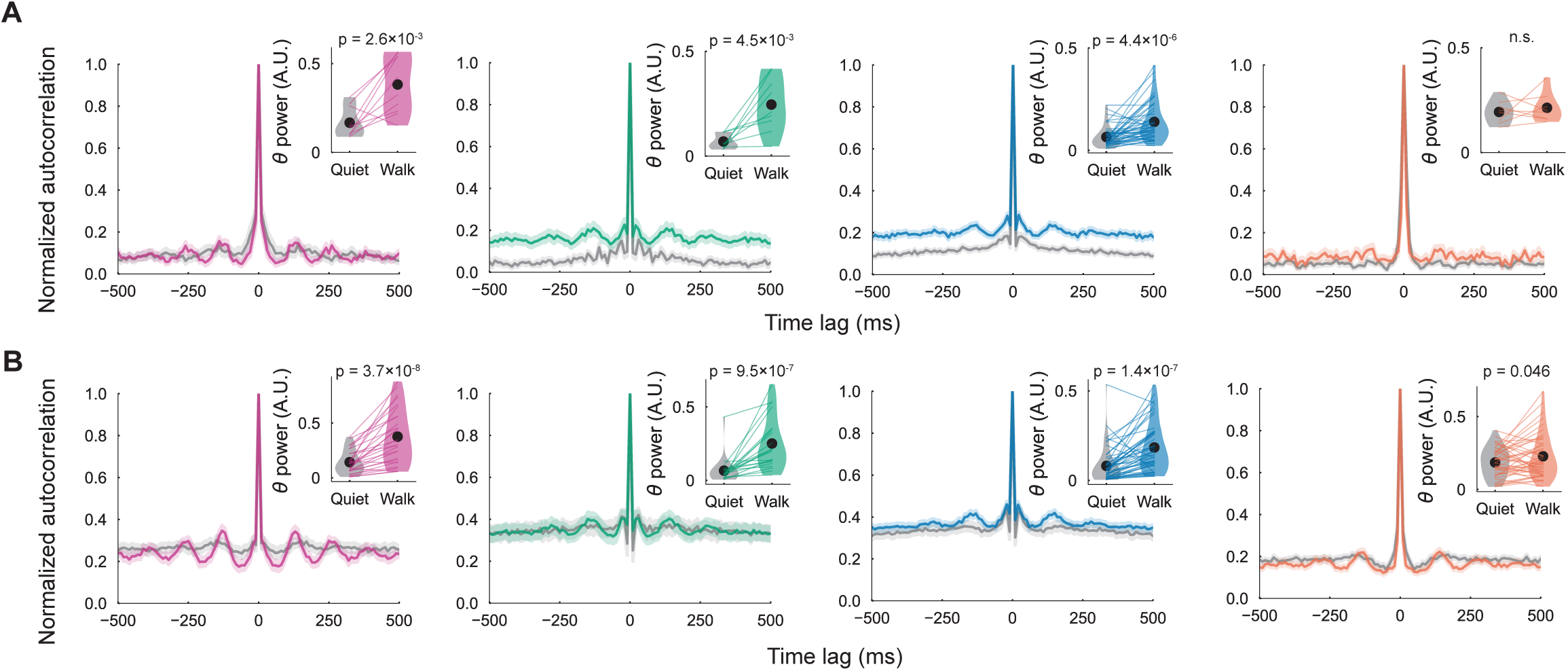
Theta modulated spiking in CA1 neurons, related to Figure 3. (A) Population averaged spike-triggered autocorrelograms during quiet (gray) and walking (colored) without optogenetic stimulation, normalized by the total number of spikes. Shaded area represents s.e.m.. All interneuron types exhibit theta-modulated firing during walking, while PCs do not. Insets: Violin plots comparing relative theta power (PSD power summed between 6-9 Hz and normalized by the total power) in the spike-triggered autocorrelograms between the two behavioral states. Only cells that fired at least 20 spikes in both behavioral states were included in the analysis. (B) Population averaged spike-triggered autocorrelograms with optogenetic stimulation, similar to (A). PCs show theta-modulated firing in both quiet and walking states, but display an additional cycle of theta entrainment during walking compared to quiet.

**Figure S4.**
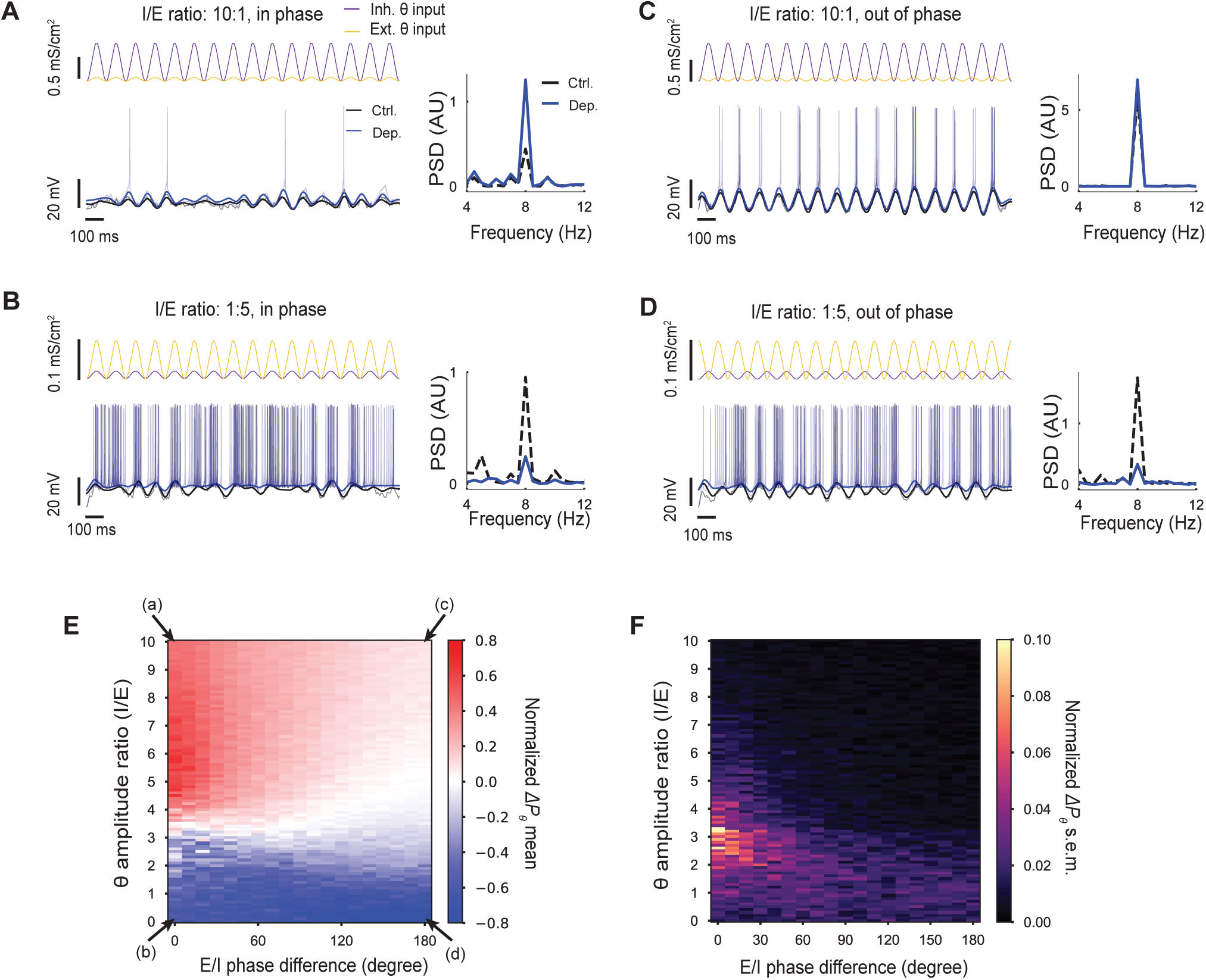
Simulations of a conductance-based model neuron receiving excitatory and inhibitory theta-oscillating inputs, related to Figure 3. (A) Example simulation with excitatory (E) and inhibitory (I) theta-oscillating conductances in phase and an I/E conductance ratio of 10. Top left, time courses of E and I conductances. Bottom left, corresponding Vm traces with and without prolonged depolarization. Right, PSDs of the simulated traces within theta band. (B-D) Same as (A), but showing simulations with different phase differences and I/E ratios. (E) Heatmap showing the mean normalized difference in theta power between control and depolarized conditions as a function of the theta amplitude ratio (I/E) and phase difference between inputs. The normalized difference in theta power is calculated as the change in theta power between the depolarized and control conditions, normalized by their combined magnitude. (F) Same as (E), but showing the s.e.m..

**Figure S5.**
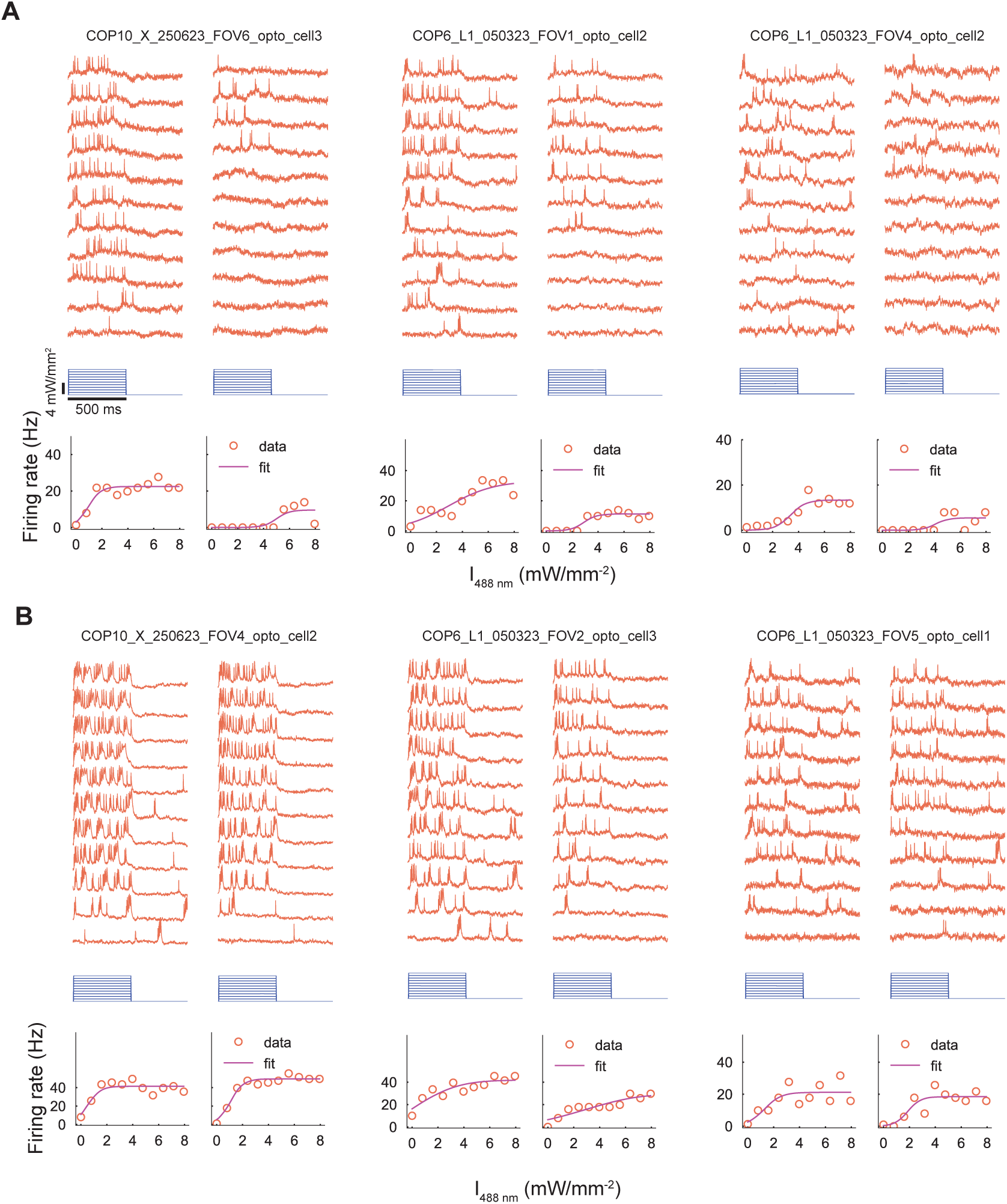
F-I curves of pyramidal cells resembling step functions during walking, related to Figure 4. (A) F-I curves of three pyramidal cells that resemble step functions during walking (a characteristic of Type II neurons). These cells are recorded from separate FOVs in two animals. (B) Other pyramidal cells do not show step function-like behavior in their F-I curves, recorded in the same imaging session as the cells in (A).

**Figure S6.**
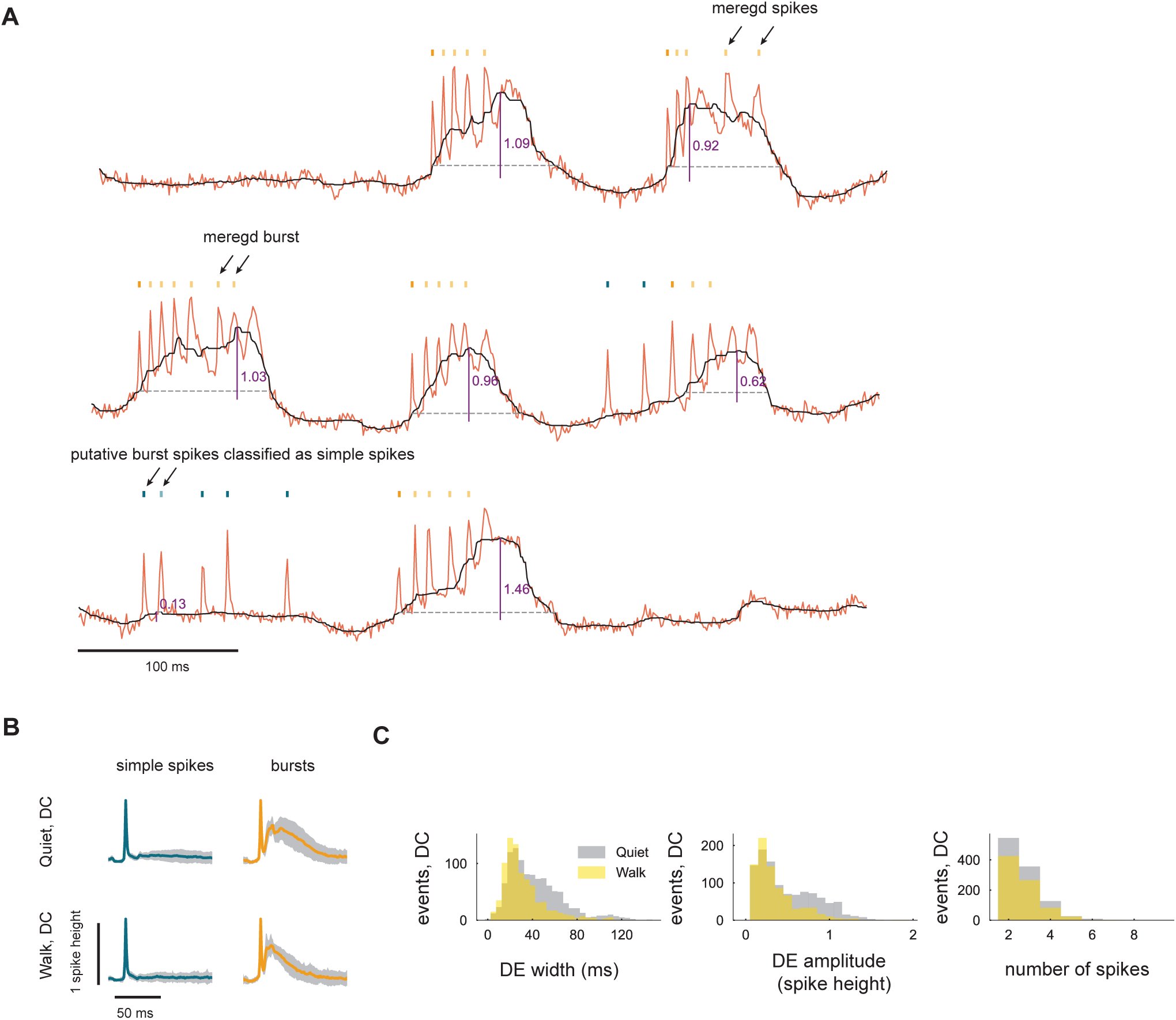
Bursts during prolonged depolarization, related to Figure 5. (A) Details of burst detection. The process begins by identifying putative bursts using an ISI threshold of 14 ms, followed by merging spikes (example in the top trace) or bursts (example in the middle trace) into bursts if they occur within a depolarization event (DE) with an amplitude larger than 0.4. Finally, burst spikes with DE amplitudes below 0.15 are reclassified as simple spikes (example in the bottom trace). Fluorescence traces (red) and subthreshold Vm (black) from three example cells are shown. Orange ticks above the traces denote burst spikes, while teal ticks denote simple spikes. Saturated ticks represent the first spike within a given burst/putative burst, while opaque ticks indicate the subsequent spikes within the same burst. Solid purple vertical lines indicate the amplitude of DEs, and black dashed lines mark the duration of DEs. (B) Population-averaged spike waveforms for simple spikes (left) and bursts (right) during prolonged depolarization across various experimental conditions. (C) Distributions of DE width, DE amplitude, and the number of spikes per burst during prolonged depolarization.

**Figure S7.**
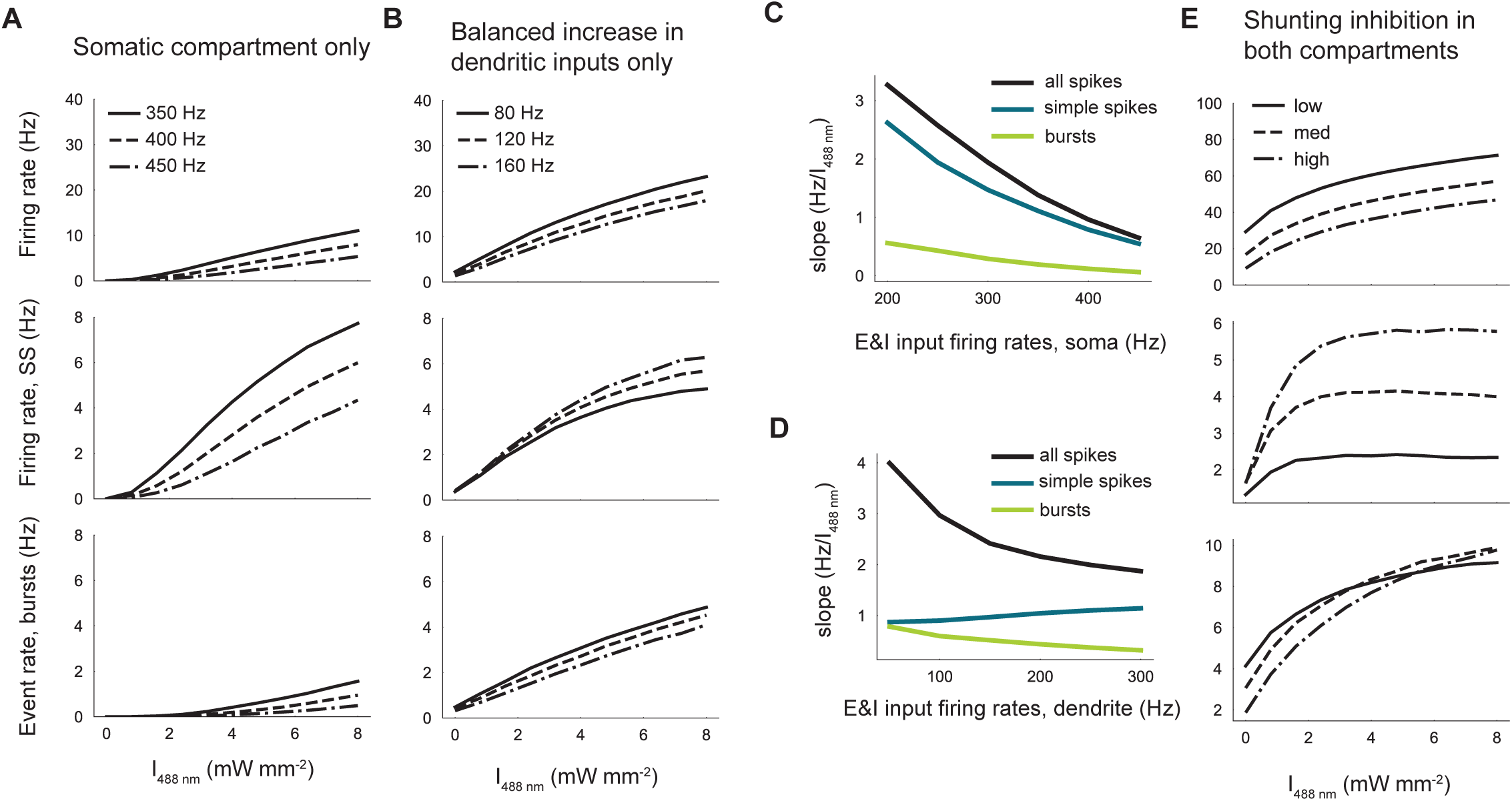
F-I curves modulated by different configurations of E/I inputs in a two-compartment model neuron, related to Figure 6. (A) F-I curves for all spikes (top), simple spikes (middle), and bursts (bottom) when the dendritic compartment is decoupled from the somatic compartment, illustrating the effects of somatic synaptic inputs alone on F-I curves. A balanced increase in synaptic input firing rate in the somatic compartment primarily modulates the gain of simple spikes. (B) F-I curves illustrating the effect of varying dendritic synaptic input frequencies while keeping somatic inputs constant (350 Hz), demonstrating dendritic contributions to gain modulation. A balanced increase in dendritic synaptic input reduces burst gain while increasing simple spike gain, effectively transforming bursts into simple spikes. (C) For neuron in (A), the F-I slopes for all spikes, simple spikes, and bursts decrease as the somatic input firing rates increase. (D) For neurons in (B), the F-I slopes for all spikes and bursts decrease with increasing dendritic input firing rates, whereas the F-I slopes for simple spikes increase. (E) F-I curves when the somatic and dendritic compartments receive increased inhibitory inputs. The excitatory synaptic input rates were fixed to 140 Hz and 80 Hz for the somatic and dendritic compartment, respectively. Low, medium, high indicate the strength of the inhibitory synaptic input rates, corresponding to somatic input frequencies of 100, 120, and 140 Hz, and dendritic input frequencies of 40, 60, and 80 Hz, respectively.

**Figure S8.**
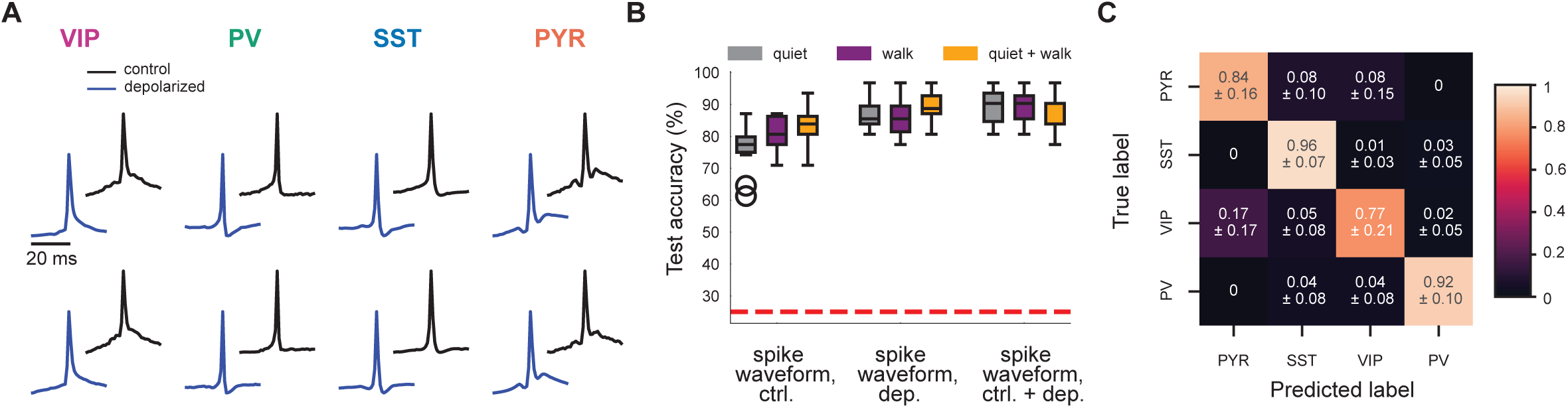
Decoding molecular identity with a 1-D CNN using shorter spike waveforms, related to Figure 7. (A) Spike waveforms (grand average) for each cell type during quiet (top) and walking (bottom) states, with (blue) and without (black) optogenetic stimulation. (B) Performance of the 1-D CNN trained on 70% of the experimental data for each cell type, evaluated across various data types, including spike waveforms from the quiet state, the walking state, and a combination of both states. Results are the mean and standard deviation of 10 random train-test splits. CNN trained with concatenated spike waveforms from control and depolarization conditions during walking showed the best overall test accuracy of 0.89 ± 0.02 (mean ± s.d.). (C) Confusion matrix of the best performing CNN, showing the accuracy of decoding each cell type (mean ± s.d.) along the diagonal and the probability of misclassification into other cell types in the off-diagonals.

